# Did Iron Suppress Eukaryote Emergence and Early Radiation?

**DOI:** 10.1101/2025.05.29.655150

**Authors:** Zachary Adam, Sarah Slotznick, Ann Bauer, Esther Stewart, Eric Roden, Shanan E. Peters, Kurt Konhauser, Betül Kaçar

## Abstract

The last eukaryotic common ancestor (LECA) is widely thought to have been an oxygen-respiring organism, arising through endosymbiosis when a free-living bacterium became the mitochondrion. Owing to the mitochondrion’s central metabolic task of oxidative phosphorylation, oxygen availability has long been a hypothesized driver of eukaryogenesis. However, this hypothesis is challenged by a temporal disconnect, spanning several hundred million years, between the earliest geochemical evidence for oxygen in the environment (∼3.2-2.5 Ga) and the oldest widely accepted eukaryotic fossils (∼1.7 Ga). Notably, the earliest candidate eukaryotes appear contemporaneous with the cessation of major iron deposits and rise of sulfide- and sulfate-rich marine sediments in coastal environments. Here, we integrate Proterozoic surface geochemical records with the microbial biochemistry of iron to examine potential environmental constraints on early eukaryotic evolution. Iron bioavailability exerts complex and often antagonistic effects on both aerobic and anaerobic microbial lineages that contributed to LECA. Elevated iron levels likely disrupted cellular homeostasis, particularly by destabilizing labile iron pools and promoting oxidative damage to bacterial lipids. The programmed cell death pathways known as ferroptosis, which is widespread among eukaryotic lineages, may trace its origins to iron-rich conditions in Archaean and Paleoproterozoic seawater and LECA. Our findings challenge oxygen-centered paradigms of eukaryogenesis and reframes the long-recognized temporal gap as a consequence of iron-mediated physiological constraints.

## Primary Endosymbiosis

Our understanding of eukaryotic origins has advanced markedly over the last decade. While eukaryotes were long considered as a distinct domain within the tree of life *(1)*, recent phylogenetic analysis of eukaryotic genomes and organelles have increasingly supported the view that eukaryotes arose through endosymbiosis of bacterial and archaeal lineages *(2)*. This interpretation is reinforced by recent discoveries of Asgard archaea, whose genome and remarkably variable cellular morphology *(3)* place Eukaryotes within the clade *(4)*. Collectively, the evidence points to a model in which eukaryotes emerged from what were free-living bacterial and archaeal progenitors. The mitochondrial progenitor was almost certainly an oxygen-respiring, heterotrophic bacterium that inhabited a near-surface environment *(5-8)*. The precise ecological and environmental drivers of this amalgamation remain unknown, but the available data suggest that the primary endosymbiotic event likely occurred during the Paleoproterozoic era (Figure 1, black band), a period marked by the rise of atmospheric oxygen, the cessation of widespread Fe-rich sediment deposition, and the appearance of the earliest credible eukaryotic microfossils.

**Figure 1.**
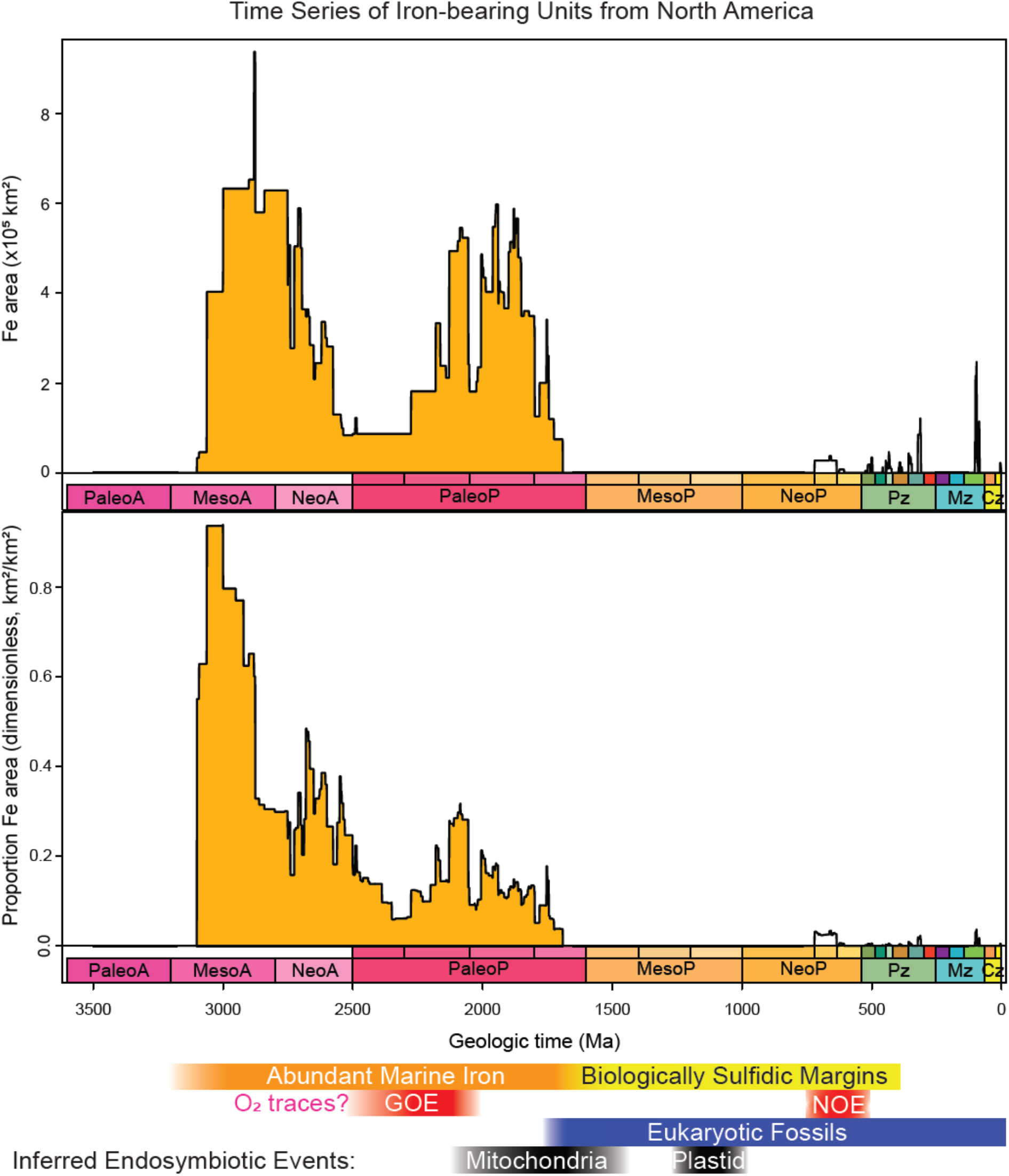
Frequency and distribution of iron-rich sediments over time: Abundance of iron-rich sedimentary units (i.e., units with “iron formation” or “ironstone” as lithologies or “ferruginous” as a lithology adjective) in the North American Macrostrat database *(30)*. Top: Aerial extent of iron-rich sediments (km^2^). Bottom: Time-varying proportion of iron-rich units vs all sedimentary and metasedimentary units (km^2^/km^2^). Oxygen events (red bars) following *(37, 42, 43)*. Timing of key eukaryote endosymbiotic events (black bars) following *(44-49)*. Timing of eukaryotic fossils (blue band) following *(50)*. Timing of biologically sulfidic margins (yellow band) following *(37, 40, 51)*.

Here we begin with a foundational and widely accepted premise: all cells have an intrinsic need to actively maintain an internal chemical environment that supports essential biochemical processes. Cellular self-regulation, known as homeostasis, involves multiple overlapping systems that enable cells to maintain stable internal conditions. Because oxygen and iron differ markedly in both their chemistry and cellular roles, each requires specialized and distinct homeostatic mechanisms *(9)*. During the Paleoproterozoic—notable for its redox fluctuations and geochemical instability—maintaining homeostasis would have posed significant challenges, particularly for organisms occupying near-surface environments. These environmental pressures likely imposed stringent constraints on both the bacterial and archaeal progenitors of eukaryotes, as well as on the earliest eukaryotic cells.

## INTRODUCTION

### Archaean and Proterozoic Redox Conditions

The surface chemical conditions of the very early Earth were likely to have included vanishingly little (if any) molecular oxygen *(10)*, but this changed with the evolution of oxygenic photosynthesis. Changes in sinks and abiotic redox balance, the origination of oxyphototrophy, and the burial of photosynthetically produced organic carbon drove oxygen levels in the atmosphere to rise over a time period ranging from about 2.4-2.1 billion years ago (Ga), a period referred to as the Great Oxidation Event or GOE (Figure 1, red band; *(11-13)*. Molecular clock estimates place the origins of basal oxyphototrophic groups just prior to the GOE *(14-16)*, and there are geochemical and geologic and geologic indications interpreted as ephemeral oxygen from units with ages ranging from 3.2-2.5 Ga (*(17)* but c.f. *(18-22)*). Regardless of exactly when oxygenic photosynthesis first emerged on Earth, what seems certain is that by around 2.5 Ga, crown group oxyphototrophic organisms had evolved and began producing oxygen as a metabolic waste product *(23)*.

The Archean oceans were anoxic and rich in dissolved ferrous iron (Fe^2+^), as evidenced by the widespread deposition of banded iron formations (BIFs)—the primary source of industrial iron ore. These uniquely Precambrian chemical sedimentary rocks typically consist of alternating layers, ranging from metre- to sub-millimetre-thick, composed of iron-rich minerals and silicate or carbonate phases. Their bulk composition generally falls within ∼20–40 wt.% Fe and ∼40–60 wt.% SiO_2_ (see *(24)*, for review). It is now widely accepted that the iron in these deposits originated as Fe(II) sourced from hydrothermal systems, although recent work also suggests some input from recycled continental sediments *(25)*. Dissolved Fe^2+^ concentrations in seawater are estimated to range from 0.03 to 1 millimolar (mM) *(26-29)*.

Figure 1 summarizes results from the Macrostrat database of North America *(30)* for any sedimentary rock unit that includes iron-related sediments in its lithological description. The rock record indicates that deposition of BIF was limited between ∼ 2.4 and 1.9 Ga *(31)*. A final widespread episode of large iron formation deposition at 1.88-1.85 Ga (predominantly as shallow-water granular iron formations, or GIFs) indicates a return to ferrous iron-rich sedimentary basins. This change may be associated with an increase in mantle plume activity *(31)*. An alternative (though not mutually exclusive) hypothesis attributes this episode to a decline in atmospheric and hence oceanic oxygen levels *(32)*. There is a termination of prominently iron-bearing units at ∼1.7 Ga. There likely remained an ongoing contribution of ferrous iron from marine hydrothermal vent systems throughout the Mesoproterozoic but these are understood not to have led to notable reservoir accumulations *(33, 34)*, at least in continental shelf settings. For these reasons, ∼1.8-1.6 Ga is considered to represent a marked transition from oxygenated shallow seawater to complexly stratified marine environments with underlying euxinic intermediate layers (Figure 1, yellow band) overlying anoxic (potentially ferruginous) deep waters *(35-41)*. This transition coincides with the earliest appearance of microfossils with eukaryotic synapomorphies (Figure 1, blue band). There appears to be no obvious correlations between rises in oxygen (red bands) and key endosymbiotic events (black bands) *(2)*.

## RESULTS AND DISCUSSION

### Early Eukaryote Biochemistry

#### Redox Homeostasis

Organisms that generate, respire or exist in proximity to oxygen must contend with its potentially damaging effects on cellular biochemistry *(52)*. It seems likely that prokaryotic (i.e., bacterial) organisms inhabiting shallow, oxygenated environments were the first to evolve the homeostatic apparatus necessary for coping with oxidative stress, mechanisms that were later inherited by stem and crown eukaryotic groups *(53)*. This apparatus, by definition, required a tight coupling between the intracellular and extracellular environments, as well as between the once-independent metabolic cycles of the bacterial and archaeal progenitors. Given that the progenitors to LECA had distinct metabolic needs, physiologies and oxygen tolerances, the challenges of maintaining cellular homeostasis would have been significant and unprecedented – perhaps so unique that homeostasis holds key insights into the overall process of endosymbiosis itself. Eukaryote homeostasis may thus provide a promising approach to bridging Paleoproterozoic environmental conditions with emerging theories of eukaryogenesis.

Oxidative stress is mitigated through a complex network of thiol (-SH)-containing compounds, with cysteine and glutathione (GSH) playing central roles in eukaryotic systems. The amino acid cysteine is found at a nexus of redox homeostasis, metabolism, translation, and cell-cycle control, functions essential to all organisms regardless of their metabolic strategy. Notably, cysteine is also the most abundant low-molecular-weight thiol found outside the cell *(54)*, highlighting its importance as an extracellular redox buffer. GSH, a tripeptide composed of L-glutamic acid, L-cysteine and glycine, is directly tied to cysteine metabolism *(55)*. GSH figures prominently in cellular homeostasis as it is the most abundant low-molecular-weight thiol redox buffer found inside eukaryotic organisms and most Gram-negative bacteria *(56-58)*. Phylogenetic studies suggest that the second step of GSH synthesis evolved in oxyphotobacteria in response to the GOE, and was subsequently disseminated to proteobacteria through horizontal gene transfer (HGT) *(59)*. This biosynthetic capability likely found its way into LECA either through mitochondrial endosymbiosis or via independent HGT events into stem-group eukaryotes *(59)*. In extant eukaryotes, GSH levels are finely regulated across different cell types and within different subcellular organelles to maximize cellular fitness *(60)*. GSH’s critical role in maintaining compartmentalized redox homeostasis suggests that the evolution of thiol-based buffering systems was a foundational step in the emergence of complex eukaryotic cells, linking redox adaptation directly to the evolutionary path of eukaryogenesis.

#### Iron toxicity and homeostasis

Proterozoic microbial communities likely thrived along continental margins, where nutrient availability (including iron) was highest *(61-63)* due to a combination of deep-water upwelling and terrestrial runoff *(64, 65)*. These dynamic coastal zones were also subject to increasing levels of dissolved oxygen and reactive oxygen species (ROS), creating challenging conditions for microbial life over hundreds of millions of years. While the effects of rising oxygen and ROS on early eukaryotes have been explored in depth (i.e.,*(53, 66, 67)*), the consequences of sustained iron deposition *(68)* and iron toxicity on early microorganisms (see *(69-75)*) remain comparatively underexamined *(76, 77)*. Moreover, iron imposes specific homeostatic demands on cells. Both the bacterial and archaeal progenitors of early eukaryotes required finely tuned redox and iron regulatory systems, but these lineages differed significantly in physiology, metabolism, and oxygenation tolerance—factors that shaped how homeostasis was achieved in the face of environmental stressors. If a causal relationship exists between the gradual decline of marine iron and the rise of eukaryotic complexity and abundance, it may be illuminated through the lens of cellular iron biochemistry. Understanding how ancient cells managed the dual challenge of oxygen and iron toxicity could offer key insights into the selective pressures that drove the emergence and global expansion of eukaryotic life.

Aqueous Fe^2+^ poses a significant biochemical hazard under aerobic conditions due to its participation in the Fenton Reaction - a reaction in which Fe^2+^ reacts with water and oxygen to form ferric iron (Fe^3+^) and highly reactive hydroxyl radicals. These radicals can trigger damaging oxidation reactions that target many different biochemical compounds in a cell *(78)*. The reactivity of Fe^2+^ can be photocatalytically accelerated in the presence of ultraviolet and visible light through photo-Fenton reactions which utilize similar radical-forming pathways and occur in both abiotic environmental settings and within cells *(79)*. Although the toxicity of Fe^2+^ in oxic environments is well documented, it is less commonly appreciated that Fe^2+^ can also be detrimental under anaerobic conditions *(70)* (Figure 2). The tolerance range for the few anaerobic organisms studied is on the order of ∼1 mM levels of Fe^2+^ *(69, 80, 81)*. These concentrations broadly correspond to the inferred range of Archaean seawater iron concentrations, though localized conditions in iron-rich sediments or restricted basins may have reached significantly higher concentrations *(28)*. In general, organisms exhibit greater iron tolerance under aerobic conditions, where Fe^3+^ predominates. Ferric iron is less soluble, less toxic and more readily sequestered or trafficked through biological pathways. Nevertheless, only a limited number of microbes have evolved specialized systems that allow them to cope with exceptionally high iron loads. Some heterotrophic bacteria in acidic environments can tolerate Fe^2+^ concentrations of up to 9 mM *(82)*, while a few specialized protists can persist in environments with measured Fe^2+^ concentrations as high as 40 mM *(83)*. These outliers underscore the extremity of such conditions, and the specialized adaptations required to survive them, suggesting that iron toxicity likely exerted persistent pressures on early microbial ecosystems, including those that gave rise to eukaryotes.

**Figure 2.**
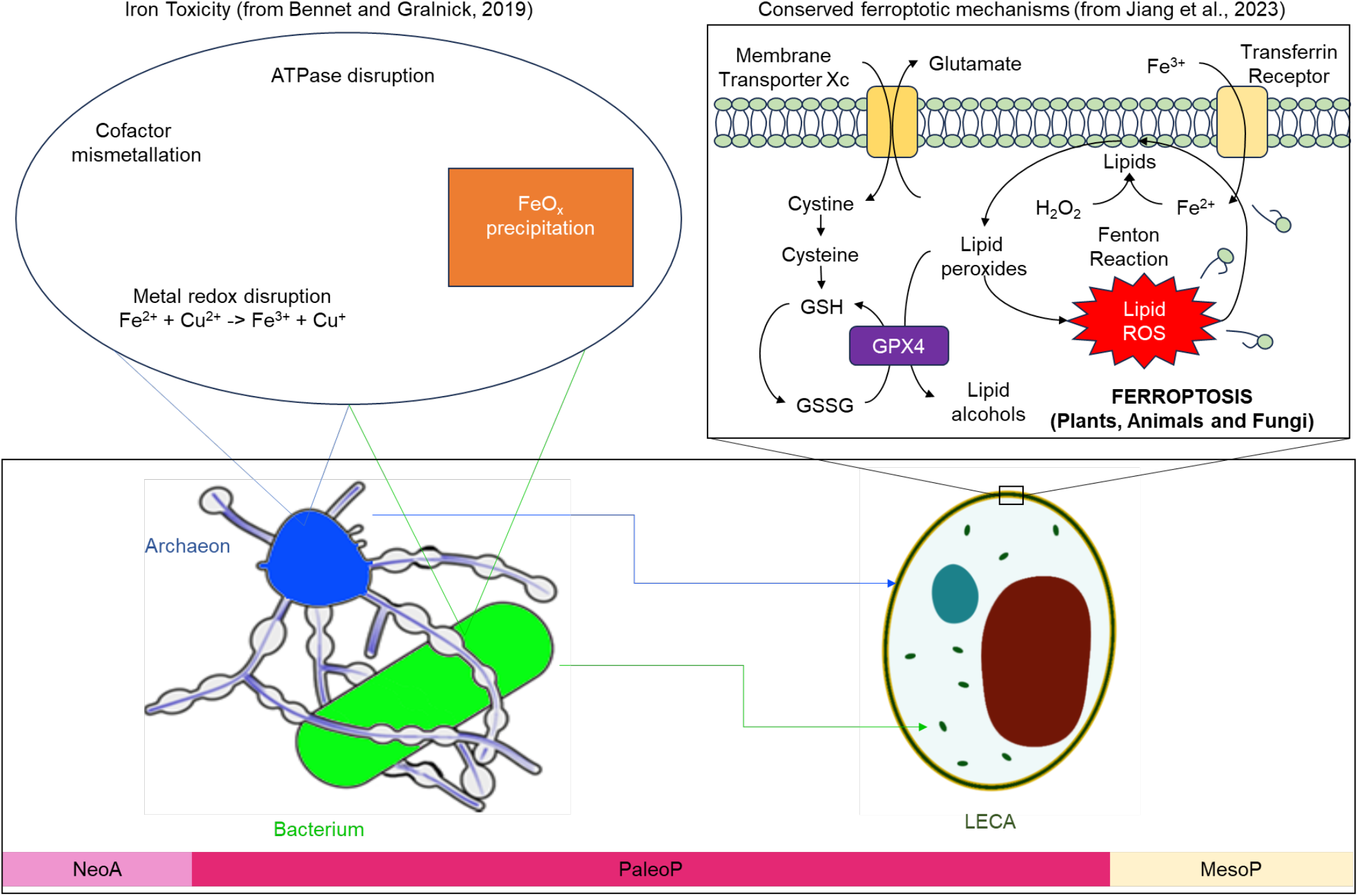
Iron disruption through endosymbiosis: An overview of disruptive effects of iron throughout endosymbiosis, from ecological associations and biochemistries of archaea and bacteria *(69)* to LECA *(90, 94)*. These effects may underlie a mechanistic connection between the cessation of marine iron deposition and the radiation of the earliest eukaryotes.

The overarching implication from biochemistry is that iron, at concentrations inferred for Archaean and Paleoproterozoic oceans based on the abundance of iron-rich sediments, may have posed a significant physiological challenge to both aerobic and anaerobic microorganisms (Figure 2). This is particularly relevant for aerobic, heterotrophic bacterial analogs of the mitochondrial progenitor, which are known to be especially susceptible to iron-induced oxidative stress *(70, 84)*. This leads us to two conclusions. The first is that we currently lack a comprehensive understanding, let alone a systematic framework, for evaluating how dissolved Fe^2+^ influenced Paleoproterozoic microbial evolution, macroevolutionary trajectories, and the endosymbiotic origins of eukaryotes. Second, it suggests that Paleoproterozoic shallow marine microorganisms and ecologies differed substantially from modern marine proxies.

The uncertainties surrounding iron toxicity and microbial ecology, while limiting in some respects, also open new possibilities for reconstructing the emergence of eukaryotes, particularly by tracing the history of iron homeostasis in prokaryotes and its importation into eukaryote physiology. The mitochondrion is appreciated for its role in aerobic respiration, but it is also a central hub of eukaryote iron metabolism *(85, 86)*. As with redox homeostasis, GSH also figures prominently in iron regulation. One noteworthy aspect of iron homeostasis is that all organisms require some minimal amount of iron; while anaerobic organisms can thrive without any oxygen, no life form can dispense entirely with iron. It is an essential metal cofactor in a wide array of enzymatic reactions, meaning that all organisms must tightly regulate intracellular iron concentrations while also adapting to external iron fluctuations *(87)*. Environmental variability of iron availability is extremely challenging for maintaining homeostasis *(88)*. Given the prolonged deposition of iron-rich sediments during the Archean and Paleoproterozoic and the noted shift to shallow water GIFs after the GOE, it is likely that iron imposed complex and shifting pressures on microbial populations inhabiting shallow marine environments—some of which ultimately became integral to early eukaryotic cell biology.

The universal need for iron, coupled with the impossibility of fully mitigating its toxic effects under both anaerobic and aerobic conditions, places complex biochemical demands upon all eukaryotic cells harboring some form of a mitochondrion *(89)*. The dual role of iron as both an essential nutrient and a potent catalyst of cellular damage has led to the evolution of sophisticated regulatory mechanisms, including a conserved form of programmed cell death known as ferroptosis *(90)*. This process is characterized by the oxidative degradation of membrane lipids in response to iron accumulation and iron-induced ROS generated via the Fenton reaction *(91)*. In ferroptosis, once initiated, the oxidation of a single lipid molecule can propagate rapidly: lipid fragments act as secondary oxidants, degrading neighboring lipids and amplifying cellular damage in a runaway cascade. If not curtailed, this feedback loop leads to cellular self-destruction. While seemingly paradoxical, the evolutionary persistence of ferroptosis across diverse eukaryotic lineages reflects its adaptive value. Under conditions of nutrient stress or environmental perturbation, the death of individual cells can release iron, amino acids, and other critical resources to nearby co-species, enhancing the resilience and survival of the larger population *(78)*.

The timing of the development of ferroptosis in eukaryotes is unknown, but mitochondrial iron metabolism regulates many of its features *(90, 92)* and there are some indications of its antiquity. Components of the ferroptotic machinery (Figure 2), including the enzymes and sensory mechanisms responsible for its regulation, are present across diverse eukaryotic clades, including animals, plants, and fungi *(93-95)*. A phylogenetic analysis of a key ferroptosis surveillance enzyme, glutathione peroxidase 4 (GPX4), indicates that divergent orthologs in plants and animals share a common ancestor with variants also present in bacteria and archaea *(96)*. Ferroptosis has not yet been observed in archaea *(90)* but a behavior consistent with ferroptosis was recently identified in the cyanobacterium *Synechocystis*; the ferroptotic mechanism acts upon lipids within the thylakoid membrane, which share a similar composition to those in eukaryotic cells *(97)*. This may link the development of this cellular process to the abundance of Paleoproterozoic iron deposits.

One fascinating implication of the antiquity of ferroptosis relates to the cryptic origins of eukaryotic lipids. Ferroptosis is closely connected to lipid metabolism *(91)*. While eukaryotic cell membranes are composed of diverse fatty acid lipid bilayers, similar to those found in bacteria, the primordial endosymbiotic host was likely an archaeon with monolayer lipids composed of isoprenoid chains *(98)*. This raises the question of what prompted a shift from archaeal to bacterial lipids, and its relative timing to eukaryogenesis. It has been suggested that the adoption of bacterial lipids provided functional advantages to the evolving eukaryotic cell such as improved membrane flexibility or permeability *(98-100)*, but iron bookends the entire process (Figure 2). Complicated iron-, oxygen- and sulfide-fluctuating environments of the early Paleoproterozoic present a novel instigating pressure for greater lipid versatility in LECA progenitors. Conserved lipid- and thiol-dependent ferroptotic mechanisms implicate iron’s biochemical antagonism through the appearance of LECA.

## CONCLUSIONS

The circumstances aligning iron deposition in Archaean and Paleoproterozoic environments, the development of oxygen and iron homeostatic machinery, and the inferred timing of mitochondrial endosymbiosis suggests that eukaryotes were borne of a complicated history of cellular homeostasis. This history involves a triumvirate of chemicals that were environmentally abundant and biochemically useful, but toxic in excess to different organisms throughout the Paleoproterozoic era: iron, oxygen and sulfide (Figure 3). Paleoproterozoic fluctuations in all of these chemicals posed tremendous challenges to homeostasis for microorganisms. Outside the cell, it is exceptionally difficult to consider any one of these chemicals in isolation as they react readily with one another (Figure 3). Within the cell, homeostatic mechanisms employ sulfide (in the form of thiols) to control harmful levels of oxygen and iron. Iron, in particular, is implicated in antagonistic biochemical roles by disrupting labile iron, enzymes, and lipids for aerobic and anaerobic eukaryotic progenitors. The origins and earliest radiation of eukaryotes were permitted by an increase in dissolved oxygen, but the history of iron’s potentially suppressive biochemical effects on endosymbiosis remain unexplored (Figure 3, dotted box). Aspects of this history ought to be recoverable through a more focused application of phylogenetics to homeostatic systems, a deeper appreciation of microbial responses to redox and iron heterogeneity, and by scrutinizing oxygen’s presumed hegemony of endosymbiosis.

**Figure 3.**
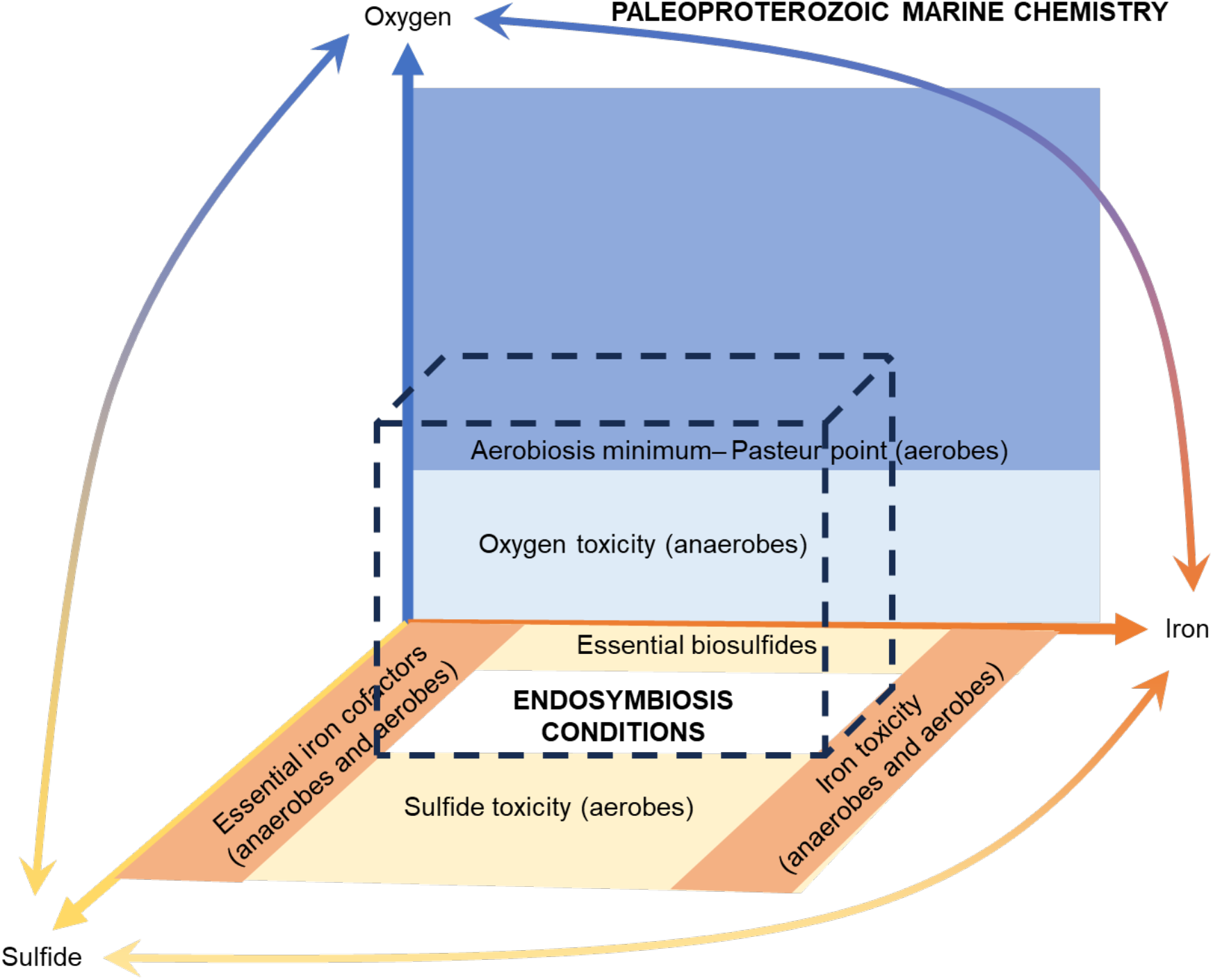
Biochemical requirements and toxicities for oxygen, iron and sulfide restrict conditions for endosymbiosis: Iron, sulfide and oxygen readily react with one another, leading to reservoir fluctuations that characterize Paleoproterozoic marine chemistry. A narrower set of interactive conditions involving these three chemicals (dotted box) are circumscribed by the context of cellular homeostatic constraints, which differ among organisms. Iron is biphasic: it is essential at lower cellular concentrations but toxic at higher cellular concentrations. Iron and oxygen homeostatic constraints must have been operable within archaea and bacteria across the process of endosymbiosis, up to and including full endogenization of the mitochondrion into a eukaryotic organelle.

Retracing the emergence of iron and redox homeostatic systems, along with the chronology of their appearance in stem and crown group eukaryotes, may offer valuable evolutionary insights into the prolonged gap between the first rise of oxygen and the earliest eukaryotes. Two distinct but not mutually exclusive possibilities warrant further study: iron directly suppressed the endosymbiotic process, or it delayed the timing of the first observable eukaryotic radiation. If the latter, then eukaryotes may have had a longer, cryptic history within low-iron refugia, such as freshwater habitats (e.g., *(101)*) that may be more poorly represented in the rock record *(102)*.

Iron’s potential role in shaping eukaryogenesis has intriguing implications for life in the universe. Atmospheric dioxygen is often considered a remotely detectable indicator of life on exoplanets, which would also underpin the presence of more complex life forms such as eukaryotes *(103, 104)*. Abundant dissolved iron ought to alter the visible appearance of oceans, and its presence or absence in conjunction with oxygen can aid the assessment of complex, eukaryote-like life beyond Earth *(105-107)*.

## METHODS

### Overview of macrostratigraphic approach

The general methodology follows that of macrostratigraphy *(30, 108, 109)*, which is based on compilations of the location, age, and properties of rock units that are lithologically- and/or geochronologically distinct at a given location. Chronologically juxtaposed rock units comprise columns, which span a geographic region that approximates rock unit extents.

### Defining Gap-Bounded Successions

In the case of sedimentary rocks, column typically consists of multiple conformable units defined by different lithologies that are juxtaposed. These units establish relative chronological control within the chronostratigraphic bin (e.g., international periods) to which the units are correlated. This group of contiguous and continuous sediments is separated from another by a temporal “gap”, establishing two “gap-bounded” successions. In the case where all sedimentary rocks are included, macrostratigraphy provided a quantitative summarization of sequence stratigraphic architecture.

### Quantifying Lithologic Attributes

Macrostratigraphic quantities, like total rock area, are calculated by aggregating all relevant units for the target lithology. In this study, we focused on units with “iron formation” or “ironstone” as lithologies or “ferruginous” as a lithology adjective. We then analyzed them over all columns in the focal area. Our study includes 949 columns spanning over 24 million square kilometers across North America.

## ACKNOWLEDGMENTS

We gratefully acknowledge support from the National Science Foundation (Sedimentary Geology and Paleobiology program, grant no. 2321014), the John Templeton Foundation (grant no 61926), NASA Interdisciplinary Consortium for Astrobiology Research (ICAR): Metal Utilization and Selection Across Eons, MUSE (grant no. 80NSSC17K0296) and the NASA Exobiology Program (grant no. NNH23ZDA001N). KK was supported by the Natural Sciences and Engineering Research Council of Canada (RGPIN-2020-05189).

## REFERENCES

1. C. R. Woese, G. E. Fox, Phylogenetic structure of the prokaryotic domain: The primary kingdoms. Proceedings of the National Academy of Sciences 74, 5088–5090 (1977).

2. D. B. Mills et al., Eukaryogenesis and oxygen in Earth history. Nature Ecology & Evolution 6, 520–532 (2022).

3. H. Imachi et al., Isolation of an archaeon at the prokaryote–eukaryote interface. Nature 577, 519–525 (2020).

4. L. Eme et al., Inference and reconstruction of the heimdallarchaeial ancestry of eukaryotes. Nature 618, 992–999 (2023).

5. F. Burki, Mitochondrial Evolution: Going, Going, Gone. Current Biology 26, R410–R412 (2016).

6. V. Hampl, A. J. Roger, in Endosymbiotic Organelle Acquisition: Solutions to the Problem of Protein Localization and Membrane Passage, S. D. Schwartzbach, P. G. Kroth, M. Oborník, Eds. (Springer International Publishing, Cham, 2024), pp. 89–121.

7. D. Speijer, How mitochondria showcase evolutionary mechanisms and the importance of oxygen. BioEssays 45, 2300013 (2023).

8. Z. Wang, M. Wu, Phylogenomic Reconstruction Indicates Mitochondrial Ancestor Was an Energy Parasite. PLOS ONE 9, e110685 (2014).

9. A. Baez, J. Shiloach, Increasing dissolved-oxygen disrupts iron homeostasis in production cultures of Escherichia coli. Antonie van Leeuwenhoek 110, 115–124 (2017).

10. T. W. Lyons, C. W. Diamond, N. J. Planavsky, C. T. Reinhard, C. Li, Oxygenation, Life, and the Planetary System during Earth’s Middle History: An Overview. Astrobiology 21, 906–923 (2021).

11. A. Bekker, in Encyclopedia of Astrobiology, M. Gargaud et al., Eds. (Springer Berlin Heidelberg, Berlin, Heidelberg, 2022), pp. 1–9.

12. W. W. Fischer, J. Hemp, J. E. Johnson, Evolution of Oxygenic Photosynthesis. Annual Review of Earth and Planetary Sciences 44, 647–683 (2016).

13. H. D. Holland, Volcanic gases, black smokers, and the great oxidation event. Geochimica et Cosmochimica Acta 66, 3811–3826 (2002).

14. J. S. Boden, K. O. Konhauser, L. J. Robbins, P. Sánchez-Baracaldo, Timing the evolution of antioxidant enzymes in cyanobacteria. Nature Communications 12, 4742 (2021).

15. C. F. Demoulin et al., Cyanobacteria evolution: Insight from the fossil record. Free Radical Biology and Medicine 140, 206–223 (2019).

16. P. Sánchez-Baracaldo, Origin of marine planktonic cyanobacteria. Scientific Reports 5, 17418 (2015).

17. A. D. Anbar et al., A Whiff of Oxygen Before the Great Oxidation Event? Science 317, 1903–1906 (2007).

18. T. Bosak, B. Liang, M. S. Sim, A. P. Petroff, Morphological record of oxygenic photosynthesis in conical stromatolites. Proceedings of the National Academy of Sciences 106, 10939–10943 (2009).

19. M. Homann, C. Heubeck, A. Airo, M. M. Tice, Morphological adaptations of 3.22 Ga-old tufted microbial mats to Archean coastal habitats (Moodies Group, Barberton Greenstone Belt, South Africa). Precambrian Research 266, 47–64 (2015).

20. N. J. Planavsky et al., Evidence for oxygenic photosynthesis half a billion years before the Great Oxidation Event. Nature Geoscience 7, 283–286 (2014).

21. A. M. Satkoski, N. J. Beukes, W. Li, B. L. Beard, C. M. Johnson, A redox-stratified ocean 3.2 billion years ago. Earth and Planetary Science Letters 430, 43–53 (2015).

22. S. P. Slotznick et al., Reexamination of 2.5-Ga “whiff” of oxygen interval points to anoxic ocean before GOE. Science Advances 8, eabj7190 (2022).

23. P. M. Shih, J. Hemp, L. M. Ward, N. J. Matzke, W. W. Fischer, Crown group Oxyphotobacteria postdate the rise of oxygen. Geobiology 15, 19–29 (2017).

24. K. O. Konhauser, A. Kappler, S. V. Lalonde, L. J. Robbins, Logan Medallist 8. Trace Elements in Iron Formation as a Window into Biogeochemical Evolution Accompanying the Oxygenation of Earth’s Atmosphere. Geoscience Canada 50, 239–258 (2023).

25. C. Wang et al., The onset of continental weathering recorded in Archean banded iron formations. Geology 53, 243–247 (2024).

26. H. D. Holland, The Oceans; A Possible Source of Iron in Iron-Formations. Economic Geology 68, 1169–1172 (1973).

27. C. Z. Jiang, N. J. Tosca, Fe(II)-carbonate precipitation kinetics and the chemistry of anoxic ferruginous seawater. Earth and Planetary Science Letters 506, 231–242 (2019).

28. J. E. Johnson, T. M. Present, J. S. Valentine, Iron: Life’s primeval transition metal. Proceedings of the National Academy of Sciences 121, e2318692121 (2024).

29. R. C. Morris, Genetic modelling for banded iron-formation of the Hamersley Group, Pilbara Craton, Western Australia. Precambrian Research 60, 243–286 (1993).

30. S. E. Peters, J. M. Husson, J. Czaplewski, Macrostrat: A Platform for Geological Data Integration and Deep-Time Earth Crust Research. Geochemistry, Geophysics, Geosystems 19, 1393–1409 (2018).

31. A. Bekker et al., Iron Formation: The Sedimentary Product of a Complex Interplay among Mantle, Tectonic, Oceanic, and Biospheric Processes*. Economic Geology 105, 467–508 (2010).

32. A. Eyster et al., A New Depositional Framework for Massive Iron Formations After the Great Oxidation Event. Geochemistry, Geophysics, Geosystems 22, e2020GC009113 (2021).

33. C. X. Liu, A. Capirala, S. L. Olson, M. F. Jansen, N. Dauphas, Ocean mixing timescale through time and implications for the origin of iron formations. Geochemical Perspectives Letters 31, 54–59 (2024).

34. C. Scott et al., Bioavailability of zinc in marine systems through time. Nature Geoscience 6, 125–128 (2013).

35. R. A. Boyle et al., Nitrogen cycle feedbacks as a control on euxinia in the mid-Proterozoic ocean. Nature Communications 4, 1533 (2013).

36. G. Luo et al., Shallow stratification prevailed for ∼1700 to ∼1300 Ma ocean: Evidence from organic carbon isotopes in the North China Craton. Earth and Planetary Science Letters 400, 219–232 (2014).

37. T. W. Lyons, C. T. Reinhard, N. J. Planavsky, The rise of oxygen in Earth’s early ocean and atmosphere. Nature 506, 307–315 (2014).

38. J. F. Slack, T. Grenne, A. Bekker, Seafloor-hydrothermal Si-Fe-Mn exhalites in the Pecos greenstone belt, New Mexico, and the redox state of ca. 1720 Ma deep seawater. Geosphere 5, 302–314 (2009).

39. J. F. Slack, T. Grenne, A. Bekker, O. J. Rouxel, P. A. Lindberg, Suboxic deep seawater in the late Paleoproterozoic: Evidence from hematitic chert and iron formation related to seafloor-hydrothermal sulfide deposits, central Arizona, USA. Earth and Planetary Science Letters 255, 243–256 (2007).

40. E. A. Sperling et al., Redox heterogeneity of subsurface waters in the Mesoproterozoic ocean. Geobiology 12, 373–386 (2014).

41. D. A. Stolper, C. B. Keller, A record of deep-ocean dissolved O2 from the oxidation state of iron in submarine basalts. Nature 553, 323–327 (2018).

42. N. J. Planavsky et al., Late Proterozoic Transitions in Climate, Oxygen, and Tectonics, and the Rise of Complex Life. The Paleontological Society Papers 21, 47–82 (2015).

43. R. G. Stockey et al., Sustained increases in atmospheric oxygen and marine productivity in the Neoproterozoic and Palaeozoic eras. Nature Geoscience 17, 667–674 (2024).

44. H. C. Betts et al., Integrated genomic and fossil evidence illuminates life’s early evolution and eukaryote origin. Nature Ecology & Evolution 2, 1556–1562 (2018).

45. L. Eme, S. C. Sharpe, M. W. Brown, A. J. Roger, On the age of eukaryotes: evaluating evidence from fossils and molecular clocks. Cold Spring Harb Perspect Biol 6, (2014).

46. T. A. Mahendrarajah et al., ATP synthase evolution on a cross-braced dated tree of life. Nature Communications 14, 7456 (2023).

47. J. Vosseberg et al., The emerging view on the origin and early evolution of eukaryotic cells. Nature 633, 295–305 (2024).

48. S. Wang, H. Luo, Dating Alphaproteobacteria evolution with eukaryotic fossils. Nature Communications 12, 3324 (2021).

49. I. Zachar, G. Boza, Endosymbiosis before eukaryotes: mitochondrial establishment in protoeukaryotes. Cellular and Molecular Life Sciences 77, 3503–3523 (2020).

50. S. M. Porter, Insights into eukaryogenesis from the fossil record. Interface Focus 10, 20190105 (2020).

51. S. W. Poulton, P. W. Fralick, D. E. Canfield, The transition to a sulphidic ocean ∼ 1.84 billion years ago. Nature 431, 173–177 (2004).

52. E. U. Hammarlund, E. Flashman, S. Mohlin, F. Licausi, Oxygen-sensing mechanisms across eukaryotic kingdoms and their roles in complex multicellularity. Science 370, (2020).

53. W. W. Fischer, J. Hemp, J. S. Valentine, How did life survive Earth’s great oxygenation? Current Opinion in Chemical Biology 31, 166–178 (2016).

54. Y.-M. Go, J. D. Chandler, D. P. Jones, The cysteine proteome. Free Radical Biology and Medicine 84, 227–245 (2015).

55. C. Cassier-Chauvat, F. Marceau, S. Farci, S. Ouchane, F. Chauvat, The Glutathione System: A Journey from Cyanobacteria to Higher Eukaryotes. Antioxidants 12, 1199 (2023).

56. R. C. Fahey, Novel Thiols of Prokaryotes. Annual Review of Microbiology 55, 333–356 (2001).

57. L. Marty et al., Arabidopsis glutathione reductase 2 is indispensable in plastids, while mitochondrial glutathione is safeguarded by additional reduction and transport systems. New Phytologist 224, 1569–1584 (2019).

58. A. J. Meyer, M. J. May, M. Fricker, Quantitative in vivo measurement of glutathione in Arabidopsis cells. The Plant Journal 27, 67–78 (2001).

59. S. D. Copley, J. K. Dhillon, Lateral gene transfer and parallel evolution in the history of glutathione biosynthesis genes. Genome Biology 3, research0025.0021 (2002).

60. M. Deponte, The Incomplete Glutathione Puzzle: Just Guessing at Numbers and Figures? Antioxidants & Redox Signaling 27, 1130–1161 (2017).

61. W. Hao et al., The kaolinite shuttle links the Great Oxidation and Lomagundi events. Nature Communications 12, 2944 (2021).

62. T. A. Laakso, D. P. Schrag, A small marine biosphere in the Proterozoic. Geobiology 17, 161–171 (2019).

63. L. A. Riedman, S. M. Porter, M. A. Lechte, A. dos Santos, G. P. Halverson, Early eukaryotic microfossils of the late Palaeoproterozoic Limbunya Group, Birrindudu Basin, northern Australia. Papers in Palaeontology 9, e1538 (2023).

64. J. Morrissey, C. Bowler, Iron utilization in marine cyanobacteria and eukaryotic algae. Front Microbiol 3, 43 (2012).

65. J. H. Ryther, D. D. Kramer, Relative Iron Requirement of Some Coastal and Offshore Plankton Algae. Ecology 42, 444–446 (1961).

66. J. Gross, D. Bhattacharya, Uniting sex and eukaryote origins in an emerging oxygenic world. Biology Direct 5, 53 (2010).

67. A. Neubeck, F. Freund, Sulfur Chemistry May Have Paved the Way for Evolution of Antioxidants. Astrobiology 20, 670–675 (2019).

68. K. O. Konhauser, A. Kappler, E. E. Roden, IRON IN MICROBIAL METABOLISMS. Elements 7, 89–93 (2011).

69. B. D. Bennett, J. A. Gralnick, Mechanisms of toxicity by and resistance to ferrous iron in anaerobic systems. Free Radical Biology and Medicine 140, 167–171 (2019).

70. J. Bird Lina, L. Coleman Maureen, K. Newman Dianne, Iron and Copper Act Synergistically To Delay Anaerobic Growth of Bacteria. Applied and Environmental Microbiology 79, 3619–3627 (2013).

71. K. Lepot et al., Iron minerals within specific microfossil morphospecies of the 1.88 Ga Gunflint Formation. Nature Communications 8, (2017).

72. E. D. Swanner et al., Modulation of oxygen production in Archaean oceans by episodes of Fe(II) toxicity. Nature Geoscience 8, 126–130 (2015).

73. D. Touati, Iron and Oxidative Stress in Bacteria. Archives of Biochemistry and Biophysics 373, 1–6 (2000).

74. C. Wang et al., Archean to early Paleoproterozoic iron formations document a transition in iron oxidation mechanisms. Geochimica et Cosmochimica Acta 343, 286–303 (2023).

75. L. M. Ward et al., Microbial diversity and iron oxidation at Okuoku-hachikurou Onsen, a Japanese hot spring analog of Precambrian iron formations. Geobiology 15, 817–835 (2017).

76. A. C. Dlouhy, C. E. Outten, in Metallomics and the Cell, L. Banci, Ed. (Springer Netherlands, Dordrecht, 2013), pp. 241–278.

77. H. G. Sherman et al., New Perspectives on Iron Uptake in Eukaryotes. Front Mol Biosci 5, 97 (2018).

78. M. S. Kwun, D. G. Lee, Ferroptosis-Like Death in Microorganisms: A Novel Programmed Cell Death Following Lipid Peroxidation. J Microbiol Biotechnol 33, 992–997 (2023).

79. R. Rani, S. Singh, in Green Chemistry and Water Remediation: Research and Applications, S. K. Sharma, Ed. (Elsevier, 2021), pp. 271–298.

80. D. Bennett Brittany, E. Redford Kaitlyn, A. Gralnick Jeffrey, Survival of Anaerobic Fe2+ Stress Requires the ClpXP Protease. Journal of Bacteriology 200, 10.1128/jb.00671-00617 (2018).

81. N. Kalantari, S. Ghaffari, Evaluation of Toxicity of Heavy Metals for Escherichia coli Growth. Iranian Journal of Environmental Health Science & Engineering 5, (2008).

82. N. Kamjunke, J. Tittel, H. Krumbeck, C. Beulker, J. Poerschmann, High Heterotrophic Bacterial Production in Acidic, Iron-Rich Mining Lakes. Microbial Ecology 49, 425–433 (2005).

83. L. A. Amaral Zettler, M. A. Messerli, A. D. Laatsch, P. J. S. Smith, M. L. Sogin, From Genes to Genomes: Beyond Biodiversity in Spain’s Rio Tinto. The Biological Bulletin 204, 205–209 (2003).

84. L. M. Ward, J. L. Kirschvink, W. W. Fischer, Timescales of Oxygenation Following the Evolution of Oxygenic Photosynthesis. Origins of Life and Evolution of Biospheres 46, 51–65 (2016).

85. B. T. Paul, D. H. Manz, F. M. Torti, S. V. Torti, Mitochondria and Iron: current questions. Expert Rev Hematol 10, 65–79 (2017).

86. D. R. Richardson et al., Mitochondrial iron trafficking and the integration of iron metabolism between the mitochondrion and cytosol. Proceedings of the National Academy of Sciences 107, 10775–10782 (2010).

87. T. Ganz, Systemic Iron Homeostasis. Physiological Reviews 93, 1721–1741 (2013).

88. P. Chandrangsu, C. Rensing, J. D. Helmann, Metal homeostasis and resistance in bacteria. Nature Reviews Microbiology 15, 338–350 (2017).

89. M. Martinez-Pastor, W. A. Lancaster, P. D. Tonner, Michael W. W. Adams, A. K. Schmid, A transcription network of interlocking positive feedback loops maintains intracellular iron balance in archaea. Nucleic Acids Research 45, 9990–10001 (2017).

90. M. Conrad et al., Regulation of lipid peroxidation and ferroptosis in diverse species. Genes Dev 32, 602–619 (2018).

91. J. Li et al., Ferroptosis: past, present and future. Cell Death & Disease 11, 88 (2020).

92. Y. e. Liu et al., The diversified role of mitochondria in ferroptosis in cancer. Cell Death & Disease 14, 519 (2023).

93. M. M. Gaschler, B. R. Stockwell, Lipid peroxidation in cell death. Biochemical and Biophysical Research Communications 482, 419–425 (2017).

94. X. Jiang, B. R. Stockwell, M. Conrad, Ferroptosis: mechanisms, biology and role in disease. Nature Reviews Molecular Cell Biology 22, 266–282 (2021).

95. L. Magtanong, P. J. Ko, S. J. Dixon, Emerging roles for lipids in non-apoptotic cell death. Cell Death & Differentiation 23, 1099–1109 (2016).

96. T. S. Trenz et al., Going Forward and Back: The Complex Evolutionary History of the GPx. Biology 10, 1165 (2021).

97. A. Aguilera et al., C-ferroptosis is an iron-dependent form of regulated cell death in cyanobacteria. Journal of Cell Biology 221, (2021).

98. G. van Meer, D. R. Voelker, G. W. Feigenson, Membrane lipids: where they are and how they behave. Nat Rev Mol Cell Biol 9, 112–124 (2008).

99. T. Gabaldón, Origin and Early Evolution of the Eukaryotic Cell. Annual Review of Microbiology 75, 631–647 (2021).

100. M. F. Siliakus, J. van der Oost, S. W. M. Kengen, Adaptations of archaeal and bacterial membranes to variations in temperature, pH and pressure. Extremophiles 21, 651–670 (2017).

101. P. Sánchez-Baracaldo, J. A. Raven, D. Pisani, A. H. Knoll, Early photosynthetic eukaryotes inhabited low-salinity habitats. Proceedings of the National Academy of Sciences 114, E7737–E7745 (2017).

102. S. E. Peters, J. M. Husson, Sediment cycling on continental and oceanic crust. Geology 45, 323–326 (2017).

103. V. S. Meadows et al., Exoplanet Biosignatures: Understanding Oxygen as a Biosignature in the Context of Its Environment. Astrobiology 18, 630–662 (2018).

104. E. W. Schwieterman, M. Leung, An Overview of Exoplanet Biosignatures. Reviews in Mineralogy and Geochemistry 90, 465–514 (2024).

105. B. Kaçar, Reconstructing Early Microbial Life. Annual Review of Microbiology 78, 463–492 (2024).

106. K. I. Rico, A. K. Garcia, M. A. Saito, B. Kaçar, A. D. Anbar, in Treatise on Geochemistry (Third edition), A. Anbar, D. Weis, Eds. (Elsevier, Oxford, 2025), pp. 337–364.

107. H. R. Rucker, B. Kaçar, Enigmatic evolution of microbial nitrogen fixation: insights from Earth’s past. Trends in Microbiology 32, 554–564 (2024).

108. Shanan E. Peters, Macrostratigraphy of North America. The Journal of Geology 114, 391–412 (2006).

109. S. E. Peters, D. P. Quinn, J. M. Husson, R. R. Gaines, Macrostratigraphy: Insights into Cyclic and Secular Evolution of the Earth-Life System. Annual Review of Earth and Planetary Sciences 50, 419–449 (2022).

