## Supplemental Information for "Did Iron Suppress Eukaryote Emergence and Early Radiation?"

### Study Design

The purpose of this study was to determine whether the presence of dissolved iron ($Fe^{2+}$ and/or $Fe^{3+}$) could have impacted the endosymbiotic process that gave rise to the Last Eukaryotic Common Ancestor (LECA). Endosymbisis is recognized as a process that spans a spectrum of relationships, ranging from loose ecological associations between archaeal and bacterial progenitors to the First Eukaryotic Common Ancestor (Archaeal-FECA- and Bacterial-FECA, respectively) to the full endogenization of the mitochondrion as an organelle (Zachar and Boza, 2020). It is extremely difficult to disentangle intermediate stages of this process based on the genomic and physiological analysis of descendants to LECA, given the long lengths of the branches that separate the beginning and end of endosymbiosis; more, well-defined external constraints on this evolutionary process are required (Mills et al. 2022). The research purpose was therefore formulated into a testable hypothesis: does the timing of the latest-occurring shallow marine iron deposits, the formation of which depends on the availability of dissolved iron in seawater, coincide with the earliest known eukaryotic microfossils?

### Improvement on Current State of the Art

Existing studies of marine iron fall into two general categories: histograms of Banded Iron Formation (BIF) occurrence over time (e.g., Klein, 2005 or Huston and Logan, 2004), or estimates about the likely bulk reservoir marine concentration of dissolved iron (Holland 1973; Morris 1993; Jiang and Tosca 2019; Johnson et al., 2024). Among these, BIF histograms are limited in their ability to inform the research question, in part because they are defined as a specific class of deposits in which iron is a significant compositional component and in part because BIF-only tabulations do not allow the abundance of this class to be assessed relative to all other preserved marine sediments. As a class, the definition of a BIF therefore fails to capture the broader range of environments (and by extension, econiches) where microbes would have been found over Earth's history in which iron may have been only a minor spatial or temporal (but nevertheless biochemically relevant) constituent. Examples include rivers, lakes or estuaries that were not strongly connected to the marine environment, which also left behind sizeable sediment deposits or notable microfossils, but that may contain only limited iron-rich strata. The research question necessitates a full assessment of microbial econiches in which iron may or may not have been found in abundance over time and an assessment of the frequency of iron-rich sediments relative to all other marine deposits.

For these reasons, the Macrostrat database (Peters et al., 2018, macrostrat.org) is well-suited for contributing data for a broader assessment of iron-rich sediments over time. The database is currently best representative of the continent of North America, which has been shown to be a useful proxy for capturing many worldwide patterns in continental sedimentation (e.g., Husson and Peters, 2017; Tasistro-Hart and Macdonald 2023). Macrostrat is comprehensive in that it attempts to characterize the basic properties of all known rocks in the upper crust of the continent and it offers relatively high temporal and spatial resolution throughout the period of time that would have recorded the effects of endosymbiosis on the planetary surface environment (namely, the Neoarchaean and onwards to the present day). By allowing the retrieval of all sedimentary/metasedimentary units known from the continent, as well as those associated with specific lithologies ('ironstone', 'iron formation') or lithology modifiers ('ferruginous'), Macrostrat permits the assessment of iron-rich sediments in both absolute and relative terms over most of Earth history. These results are, therefore, comprehensive and are reflective of as many microbially-dominated econiches as possible for the time range of study.

**Macrostrat Data and Availability**

All geologic column data in the Macrostrat database are divided into "projects" that generally correspond to geographic regions and coverage focus. Here, we use data from "project_id=1", which corresponds to the comprehensive dataset for North America (Peters et al., 2018, 2022). A total of 21,784 sedimentary/metasedimentary rock units (generally lithostratigraphic units, such as formations) from 949 columns, and spanning the time interval 3,960 Ma to 0 Ma, are currently in this North American dataset in Macrostrat. Of these units, 108 have “iron formation” and/or “ironstone” as a lithology and 17 have “ferruginous” as a lithology modifier.

Here, we tabulate the area covered by all iron-bearing (as well as all other) sediments/metasediments by summing the areas of the columns containing the appropriate units. However, there are other metrics, such as the proportion of all columns or proportion of all units, that yield comparable results. All of the data used herein and described above are readily accessible via the Macrostrat Application Programming Interface (API), which can be parameterized to retrieve the above units and any other arbitrary combination of them based on their full complement of properties (Peters et al. 2018; Quinn et al. 2024). Specifically relevant to our study, the following API call (URL) identifies all units containing any iron-rich lithology:

https://macrostrat.org/api/units?lith=iron%20formation,ironstone&project_id=1&response=long

A separate API call (URL) identifies all units with “ferruginous” as a lithology modifier:

https://macrostrat.org/api/units?lith_att=ferruginous&project_id=1&response=long

For the convenience of readers of this manuscript, and to enable reproducibility of the figures herein, we also include tables of the iron sediment-bearing units that are the focus of this study. Please refer to the Macrostrat database for up-to-date information about these and all other units.

**SI REFERENCES**

Holland, Heinrich D. "The oceans; a possible source of iron in iron-formations." *Economic Geology* 68, no. 7 (1973): 1169-1172.

Husson, Jon M., and Shanan E. Peters. "Atmospheric oxygenation driven by unsteady growth of the continental sedimentary reservoir." *Earth and Planetary Science Letters* 460 (2017): 68-75.

Huston, David L., and Graham A. Logan. "Barite, BIFs and bugs: evidence for the evolution of the Earth's early hydrosphere." *Earth and Planetary Science Letters* 220, no. 1-2 (2004): 41-55.

Jiang, Clancy Zhijian, and Nicholas J. Tosca. "Fe (II)-carbonate precipitation kinetics and the chemistry of anoxic ferruginous seawater." *Earth and Planetary Science Letters* 506 (2019): 231-242.

Johnson, Jena E., Theodore M. Present, and Joan Selverstone Valentine. "Iron: Life's primeval transition metal." *Proceedings of the National Academy of Sciences* 121, no. 38 (2024): e2318692121.

Klein, Cornelis. "Some Precambrian banded iron-formations (BIFs) from around the world: Their age, geologic setting, mineralogy, metamorphism, geochemistry, and origins." American Mineralogist 90, no. 10 (2005): 1473-1499.

Mills, Daniel B., Richard A. Boyle, Stuart J. Daines, Erik A. Sperling, Davide Pisani, Philip CJ Donoghue, and Timothy M. Lenton. "Eukaryogenesis and oxygen in Earth history." *Nature ecology & evolution* 6, no. 5 (2022): 520-532.

Morris, R. C. "Genetic modelling for banded iron-formation of the Hamersley Group, Pilbara Craton, Western Australia." *Precambrian Research* 60, no. 1-4 (1993): 243-286.

Peters, Shanan E., Jon M. Husson, and John Czaplewski. "Macrostrat: a platform for geological data integration and deep-time earth crust research." *Geochemistry, Geophysics, Geosystems* 19, no. 4 (2018): 1393-1409.

Peters, Shanan E., Daven P. Quinn, Jon M. Husson, and Robert R. Gaines. "Macrostratigraphy: insights into cyclic and secular evolution of the Earth-life system." *Annual Review of Earth and Planetary Sciences* 50, no. 1 (2022): 419-449.

Quinn, Daven P., Casey R. Idzikowski, and Shanan E. Peters. "Building a multi-scale, collaborative, and time-integrated digital crust: The next stage of the Macrostrat data system." *Geoscience Data Journal* 11, no. 1 (2024): 11-26.

Tasistro-Hart, Adrian R., and Francis A. Macdonald. "Phanerozoic flooding of North America and the great unconformity." *Proceedings of the National Academy of Sciences* 120, no. 37 (2023): e2309084120.

Zachar, István, and Gergely Boza. "Endosymbiosis before eukaryotes: mitochondrial establishment in protoeukaryotes." Cellular and Molecular Life Sciences 77, no. 18 (2020): 3503-3523.
