## Supplemental Table 1 for "Did Iron Suppress Eukaryote Emergence and Early Radiation?"

SI_CSV_File_1_Ironstone_Ironformation_units

| unit_id | section_id | col_id | project_id | col_area | unit_name | strat_name_id | Mbr | Fm | Gp | SGp | t_age | b_age | max_thick | min_thick | outcrop | pbdb_collections | pbdb_occurrences |
|---|---|---|---|---|---|---|---|---|---|---|---|---|---|---|---|---|---|
| 13416 | 3536 | 454 | 1 | 10349.959 | Unnamed | | | | | | 1.9379 | 5.333 | 5 | 0 | surface | 0 | 0 |
| 37328 | 9562 | 1627 | 1 | 78237.617 | Badheart Fm | 4812 | | Badheart | Smoky | | 84.275 | 86.3 | 50 | 50 | | 0 | 0 |
| 37146 | 9519 | 1623 | 1 | 12569.534 | Badheart Fm | 4812 | | Badheart | Smoky | | 84.5 | 86.3 | 30 | 30 | | 1 | 17 |
| 37278 | 9549 | 1626 | 1 | 23715.732 | Badheart Fm | 4812 | | Badheart | Smoky | | 85.625 | 86.3 | 30 | 30 | | 0 | 0 |
| 3037 | 600 | 81 | 1 | 16088.815 | Windrow Fm | 2311 | | Windrow | | | 93.9 | 95.55 | 15 | 1 | both | 0 | 0 |
| 5237 | 1233 | 149 | 1 | 30975.908 | Corwin Fm of Nanushuk Gp | 7319 | | Corwin | Nanushuk | | 95 | 96.1 | 2700 | 1000 | both | 26 | 131 |
| 9497 | 2384 | 340 | 1 | 21013.658 | Belle Fourche Fm | 142 | | Belle Fourche | Colorado | | 95.55 | 100.5 | 75 | 0 | subsurface | 0 | 0 |
| 9596 | 2411 | 341 | 1 | 39055.152 | Belle Fourche Fm | 142 | | Belle Fourche | Colorado | | 95.55 | 100.5 | 60 | 0 | subsurface | 0 | 0 |
| 9627 | 2422 | 342 | 1 | 36781.879 | Belle Fourche Fm | 142 | | Belle Fourche | Colorado | | 95.55 | 100.5 | 50 | 0 | subsurface | 0 | 0 |
| 9713 | 2442 | 343 | 1 | 22995.643 | Belle Fourche Fm | 142 | | Belle Fourche | Colorado | | 95.55 | 100.5 | 55 | 0 | subsurface | 0 | 0 |
| 9795 | 2468 | 346 | 1 | 26650.545 | Belle Fourche Fm | 142 | | Belle Fourche | Colorado | | 95.55 | 100.5 | 73 | 0 | both | 0 | 0 |
| 9857 | 2487 | 347 | 1 | 27250.605 | Belle Fourche Fm and Mowry Fm | 142 | | Belle Fourche | Colorado | | 95.55 | 102.5834 | 260 | 0 | both | 1 | 7 |
| 9902 | 2503 | 348 | 1 | 42115.59 | Belle Fourche Fm and Mowry Fm | 142 | | Belle Fourche | Colorado | | 95.55 | 102.5834 | 85 | 0 | both | 0 | 0 |
| 11396 | 3069 | 429 | 1 | 33887.039 | Saginaw Fm | 1788 | | Saginaw | | | 310 | 322.15 | 73 | 0 | both | 0 | 0 |
| 11419 | 3079 | 430 | 1 | 26098.557 | Saginaw Fm | 1788 | | Saginaw | | | 310 | 322.15 | 50 | 20 | both | 2 | 424 |
| 11486 | 3093 | 431 | 1 | 25142.66 | Saginaw Fm | 1788 | | Saginaw | | | 310 | 322.15 | 40 | 0 | both | 2 | 52 |
| 15284 | 4051 | 497 | 1 | 36189.281 | Atoka Gp | 79 | | Atoka | | | 312 | 313.75 | 45 | 30 | subsurface | 0 | 0 |
| 12236 | 3265 | 438 | 1 | 14293.066 | New Providence Shale | 4164 | | New Providence Shale | Borden | | 349.5 | 350.5 | 85 | 65 | both | 0 | 0 |
| 12498 | 3336 | 443 | 1 | 12679.619 | Olentangy Shale | 4175 | | Olentangy Shale | | | 383.5334 | 384.3667 | 18 | 8 | both | 0 | 0 |
| 34907 | 8782 | 1513 | 1 | 11711.186 | Bird Fiord Fm | 4898 | | Bird Fiord | | | 389.5666 | 396.875 | 400 | 400 | | 8 | 28 |
| 8236 | 2070 | 307 | 1 | 11732.15 | Buttermilk Falls Limestone | 3509 | | Buttermilk Falls Limestone | | | 389.66 | 393.3 | 12 | 0 | surface | 0 | 0 |
| 273 | 66 | 13 | 1 | 6650.055 | Rockwood Fm | 1760 | | Rockwood | | | 423.7333 | 443.8 | 244 | 61 | both | 0 | 0 |
| 432 | 75 | 16 | 1 | 2824.982 | Rockwood Fm | 1760 | | Rockwood | | | 423.7333 | 443.8 | 168 | 107 | both | 0 | 0 |
| 274 | 66 | 13 | 1 | 6650.055 | Clinton Fm | 410 | | Clinton | | | 424.4667 | 438.5 | 99 | 0 | both | 0 | 0 |
| 433 | 75 | 16 | 1 | 2824.982 | Clinton Fm | 410 | | Clinton | | | 424.4667 | 438.5 | 99 | 0 | both | 1 | 29 |
| 1434 | 180 | 41 | 1 | 5738.12 | Lower Hematitic Mbr of Mifflintown Fm | 1290 | | Mifflintown | | | 428.9 | 433.4 | 3 | 0 | surface | 0 | 0 |
| 8503 | 2124 | 312 | 1 | 24061.959 | Clinton Fm | 410 | | Clinton | | | 431.9 | 434.675 | 0 | 0 | both | 24 | 465 |
| 12393 | 3305 | 441 | 1 | 13055.024 | Dayton Fm | 500 | | Dayton | | | 433.4 | 434.675 | 6 | 0 | subsurface | 0 | 0 |
| 1993 | 308 | 47 | 1 | 4248.86 | Cresaptown Sandstone Mbr | 2816 | Cresaptown Sandstone | Rose Hill | | | 436.8 | 437.65 | 15 | 0 | both | 0 | 0 |
| 1223 | 163 | 32 | 1 | 10739.39 | Cacapon Sandstone Mbr of Rose Hill Fm | 6959 | Cacapon Sandstone | Rose Hill | | | 437.225 | 437.65 | 17 | 0 | subsurface | 0 | 0 |
| 1323 | 173 | 39 | 1 | 8320.473 | Cacapon Sandstone Mbr of Rose Hill Fm | 6959 | Cacapon Sandstone | Rose Hill | | | 437.225 | 437.65 | 33 | 0 | | 0 | 0 |
| 1713 | 218 | 46 | 1 | 9995.404 | Cacapon Sandstone Mbr of Rose Hill Fm | 6959 | Cacapon Sandstone | Rose Hill | | | 437.225 | 437.65 | 30 | 0 | subsurface | 0 | 0 |
| 13327 | 3497 | 452 | 1 | 12938.724 | Brassfield Fm | 236 | | Brassfield | | | 438.5 | 440.8 | 20 | 0 | both | 0 | 0 |
| 11216 | 3034 | 31 | 1 | 27548.84 | Neda Fm | 1404 | | Neda | Maquoketa | | 445.8333 | 446.4667 | 15 | 0 | both | 0 | 0 |
| 2965 | 565 | 79 | 1 | 12598.981 | Glenwood Mbr of St Peter Fm | 7026 | Glenwood | St Peter Sandstone | | | 457.925 | 458.4 | 8 | 1 | subsurface | 0 | 0 |
| 2960 | 564 | 79 | 1 | 12598.981 | Mt Simon Sandstone | 3358 | | Mount Simon Sandstone | Dresbach | | 497.85 | 501 | 15 | 12 | subsurface | 0 | 0 |
| 3006 | 585 | 80 | 1 | 21693.297 | Mt Simon Sandstone | 3358 | | Mount Simon Sandstone | Dresbach | | 497.85 | 501 | 35 | 25 | subsurface | 0 | 0 |
| 42926 | 12151 | 308 | 1 | 2982.175 | Chestnut Hill | 61414 | | Chestnut Hill | | | 606.14 | 625.38 | 300 | 0 | | 0 | 0 |
| 42931 | 12156 | 309 | 1 | 4577.862 | Chestnut Hill | 61414 | | Chestnut Hill | | | 606.14 | 625.38 | 200 | 0 | | 0 | 0 |
| 13873 | 13553 | 464 | 1 | 27369.797 | Kingston Peak Fm | 1048 | | Kingston Peak | Pahrump | | 635 | 720 | 3000 | 0 | | 0 | 0 |
| 34208 | 8496 | 1481 | 1 | 10271.97 | Sayunei Fm | 9580 | | Sayunei | Rapitan | Windermere | 656.25 | 663.3331 | 40 | 40 | | 0 | 0 |
| 22043 | 6054 | 797 | 1 | 74726.438 | Wabush Lake | 6100 | | Wabush Lake | Gagnon | | 1690 | 1780 | 0 | 0 | | 0 | 0 |
| 14935 | 3974 | 488 | 1 | 44986.121 | Proterozoic Complexes | | | | | | 1724 | 1750 | 1500 | 0 | | 0 | 0 |
| 22290 | 6191 | 772 | 1 | 140732.812 | Kipalu Fm of Belcher Gp | 5439 | | Kipalu | Belcher | | 1744 | 1753 | 150 | 0 | | 0 | 0 |
| 22139 | 6107 | 789 | 1 | 125552.977 | Temiscamie | 6022 | | Temiscamie | Mistassini | | 1750.003 | 1899.997 | 0 | 0 | | 0 | 0 |
| 22303 | 6194 | 771 | 1 | 197740.25 | Nastapoka | 7880 | | | Nastapoka | | 1798 | 1870 | 762 | 0 | | 0 | 0 |
| 43003 | 12200 | 179 | 1 | 25895.09 | Animikie | 2366 | | | Animikie | | 1800 | 1850 | 2000 | 0 | | 0 | 0 |
| 33358 | 1499 | 196 | 1 | 11090.265 | Stambaugh Fm of Paint River Gp | 9410 | | Stambaugh | Paint River | Marquette Range | 1820.5 | 1840.75 | 30 | 25 | | 0 | 0 |
| 5943 | 1410 | 180 | 1 | 31876.141 | Emily Iron Formation Mbr | 2839 | Emily | Rabbit Lake | Animikie | | 1850 | 1862.5 | 300 | 60 | subsurface | 0 | 0 |
| 22346 | 6213 | 767 | 1 | 87516.531 | Wasekwan | 6108 | | | Wasekwan | | 1850 | 1880 | 1200 | 1200 | | 0 | 0 |
| 43002 | 12200 | 179 | 1 | 25895.09 | North Range | 72464 | | | North Range | | 1850 | 1925 | 2000 | 0 | | 0 | 0 |
| 6243 | 1499 | 196 | 1 | 11090.265 | Riverton Iron Fm of Paint River Gp | 9408 | | Riverton Iron | Paint River | Marquette Range | 1852 | 1870 | 245 | 45 | | 0 | 0 |
| 5944 | 1410 | 180 | 1 | 31876.141 | Trommald Fm | 2134 | | Trommald | Animikie | | 1862.5 | 1875 | 100 | 0 | subsurface | 0 | 0 |
| 5965 | 1415 | 182 | 1 | 22368.387 | Mbr of Virginia Fm | 2212 | | Virginia | Animikie | | 1862.5 | 1893.75 | 20 | 0 | subsurface | 0 | 0 |
| 5972 | 1417 | 183 | 1 | 23246.199 | Gunflint Iron Fm | 814 | | Gunflint Iron | Animikie | | 1862.5 | 1877.5 | 90 | 0 | both | 0 | 0 |
| 22578 | 6280 | 783 | 1 | 41036.914 | Gunflint Iron Fm | 814 | | Gunflint Iron | Animikie | | 1862.5 | 1877.5 | 183 | 0 | | 0 | 0 |
| 22390 | 6236 | 761 | 1 | 175195.859 | Unnamed | 7854 | | | Hurwitz | | 1870 | 1960 | 0 | 660 | | 0 | 0 |
| 48888 | 12554 | 577 | 1 | 151562.266 | Ketyet River Gp | 108116 | | | Ketyet River | | 1877.5 | 2180 | 4200 | 2500 | | 0 | 0 |
| 6171 | 1477 | 192 | 1 | 13468.912 | Ironwood Iron Fm | 960 | | Ironwood Iron | Menominee | Marquette Range | 1881.25 | 1937.5 | 315 | 160 | both | 0 | 0 |
| 5988 | 1422 | 184 | 1 | 27698.447 | Trommald Fm | 2134 | | Trommald | Animikie | | 1925 | 1950 | 100 | 0 | subsurface | 0 | 0 |
| 5967 | 1415 | 182 | 1 | 22368.387 | Biwabik Iron Fm | 178 | | Biwabik Iron | Animikie | | 1932.8125 | 1948.4375 | 225 | 60 | both | 0 | 0 |
| 22318 | 6201 | 770 | 1 | 220768.5 | Sokoman Fm of Knob Lake Gp | 5937 | | Sokoman | Knob Lake | Kaniapiskau | 1939.003 | 1971.997 | 0 | 722 | | 0 | 0 |
| 6192 | 1484 | 193 | 1 | 11111.584 | Magnetic Strata Near Watersmeet | | | | | | 1960 | 2005 | 2100 | 1200 | subsurface | 0 | 0 |
| 6202 | 1489 | 194 | 1 | 8274.834 | Amasa Fm | 31 | | Amasa | Menominee | Marquette Range | 1960 | 2005 | 215 | 0 | subsurface | 0 | 0 |
| 6237 | 1498 | 196 | 1 | 11090.265 | Vulcan Iron Fm | 2214 | | Vulcan Iron | Menominee | Marquette Range | 1960 | 2005 | 100 | 50 | both | 0 | 0 |
| 22319 | 6201 | 770 | 1 | 220768.5 | Ruth | 5860 | | Ruth | Knob Lake | Kaniapiskau | 1971.997 | 2005 | 0 | 722 | | 0 | 0 |
| 5949 | 1410 | 180 | 1 | 31876.141 | Randall Fm | 1698 | | Randall | Mille Lacs | | 1987.5 | 2018.75 | 0 | 0 | both | 0 | 0 |
| 6211 | 1492 | 195 | 1 | 22234.158 | Michigamme Fm | 1285 | | Michigamme | Baraga | Marquette Range | 1996 | 2023 | 150 | 0 | both | 0 | 0 |
| 42714 | 11974 | 332 | 1 | 29450.566 | unnamed shale and Fe formation | | | | | | 2000 | 2087.5 | 1000 | 0 | | 0 | 0 |
| 9541 | 2394 | 340 | 1 | 21013.658 | Arc rocks | | | | | | 2050 | 2500 | 1500 | 0 | subsurface | 0 | 0 |
| 9542 | 2395 | 341 | 1 | 39055.152 | Arc rocks | | | | | | 2050 | 2500 | 0 | 0 | subsurface | 0 | 0 |
| 22331 | 6204 | 769 | 1 | 282089.438 | Unnamed | 7872 | | | Povungnituk | | 2054.5 | 2128.75 | 0 | 0 | | 0 | 0 |
| 6214 | 1492 | 195 | 1 | 22234.158 | Negaunee Iron Fm | 1408 | | Negaunee Iron | Menominee | Marquette Range | 2077 | 2095 | 1000 | 100 | both | 0 | 0 |
| 42720 | 11979 | 346 | 1 | 26650.545 | Arc-related rocks | | | | | | 2140 | 2500 | 1000 | 0 | | 0 | 0 |
| 22339 | 6210 | 768 | 1 | 94827.734 | Unnamed | | | | | | 2162.5 | 2275 | 0 | 0 | | 0 | 0 |
| 41903 | 11630 | 512 | 1 | 35648.656 | Benchmark Iron | 69050 | | Benchmark Iron | Nemo | | 2484 | 2490 | 70 | 60 | | 0 | 0 |
| 42891 | 12128 | 1675 | 1 | 83662.969 | supracrustal Archean | | | | | | 2500 | 2620 | 0 | 0 | subsurface | 0 | 0 |
| 16020 | 4235 | 486 | 1 | 15866.498 | Wyoming Province | | | | | | 2536 | 3100 | 0 | 0 | | 0 | 0 |
| 15980 | 4225 | 514 | 1 | 12729.511 | Unnamed | | | | | | 2542 | 3092 | 0 | 0 | | 0 | 0 |
| 15811 | 4186 | 510 | 1 | 17355.143 | Wyoming Province | | | | | | 2545 | 3100 | 0 | 0 | | 0 | 0 |
| 2951 | 560 | 33 | 1 | 8337.878 | Matlock Iron Fm | 1242 | | Matlock Iron | | | 2575 | 2620 | 50 | 0 | subsurface | 0 | 0 |
| 18214 | 4664 | 576 | 1 | 143246.766 | Unnamed | | | | | | 2575 | 2725 | 0 | 0 | | 0 | 0 |
| 42730 | 11987 | 506 | 1 | 19214.545 | Wyoming Prov. Sediments | | | | | | 2599.999 | 3000 | 1000 | 0 | | 0 | 0 |
| 41900 | 11629 | 512 | 1 | 35648.656 | unnamed | | | | | | 2605 | 2644 | 1000 | 600 | | 0 | 0 |
| 21985 | 6019 | 756 | 1 | 75979.172 | Wilson Island | 7850 | | | Wilson Island | | 2650 | 2680 | 8000 | 0 | | 0 | 0 |
| 5962 | 1414 | 182 | 1 | 22368.387 | Soudan Iron Fm | 1953 | Soudan Iron Formation | Ely Greenstone | | | 2665 | 2680 | 500 | 0 | both | 0 | 0 |
| 22575 | 6278 | 783 | 1 | 41036.914 | UNNAMED | | | | | | 2665 | 2740 | 0 | 0 | | 0 | 0 |
| 22175 | 6130 | 786 | 1 | 16276.36 | Timiskaming Gp sediments | 7898 | | | Timiskaming | | 2670.01 | 2676.67 | 0 | 0 | | 0 | 0 |
| 5963 | 1414 | 182 | 1 | 22368.387 | Ely Greenstone | 5173 | | Ely Greenstone | | | 2680 | 2740 | 6000 | 3650 | both | 0 | 0 |
| 22230 | 6163 | 779 | 1 | 28548.439 | Unnamed | | | | | | 2680 | 2740 | 0 | 0 | | 0 | 0 |
| 22259 | 6177 | 775 | 1 | 63762.832 | Oxford Lake unnamed 2 | 7889 | | | Oxford Lake | | 2680 | 2710 | 0 | 0 | | 0 | 0 |
| 22096 | 6084 | 793 | 1 | 18787.191 | Unnamed | | | | | | 2695 | 2740 | 0 | 0 | | 0 | 0 |
| 22210 | 6154 | 782 | 1 | 117460.828 | Jolliffe Ore Zone in Steeprock Gp | 10529 | | Jolliffe Ore Zone | Steep Rock | | 2695 | 2710 | 400 | 100 | | 0 | 0 |
| 22280 | 6185 | 774 | 1 | 88428.953 | Unnamed | | | | | | 2700.001 | 2749.999 | 0 | 0 | | 0 | 0 |
| 22135 | 6104 | 790 | 1 | 83744.773 | Unnamed | | | | | | 2710 | 2725 | 0 | 0 | | 0 | 0 |
| 22227 | 6161 | 780 | 1 | 12891.523 | Keewatin | 2520 | | | Keewatin | | 2710 | 2880 | 0 | 0 | | 0 | 0 |
| 22214 | 6154 | 782 | 1 | 117460.828 | Keewatin | 2520 | | | Keewatin | | 2740 | 2880 | 0 | 0 | | 0 | 0 |
| 22236 | 6165 | 778 | 1 | 109073.352 | Unnamed 2 | | | | | | 2740 | 2840 | 0 | 0 | | 0 | 0 |
| 22176 | 6131 | 786 | 1 | 16276.36 | Keewatin Gp | 2520 | | | Keewatin | | 2740 | 2880 | 0 | 0 | | 0 | 0 |
| 22183 | 6136 | 785 | 1 | 27988.043 | Keewatin Group | 2520 | | | Keewatin | | 2740 | 2840 | 0 | 0 | | 0 | 0 |
| 22231 | 6163 | 779 | 1 | 28548.439 | Keewatin | 2520 | | | Keewatin | | 2740 | 2880 | 0 | 0 | | 0 | 0 |
| 22574 | 6278 | 783 | 1 | 41036.914 | KEEWATIN | 2520 | | | Keewatin | | 2740 | 2880 | 0 | 0 | | 0 | 0 |
| 22402 | 6245 | 760 | 1 | 210314.859 | Unnamed | | | | | | 2750.005 | 2999.995 | 0 | 0 | | 0 | 0 |
| 22185 | 6136 | 785 | 1 | 27988.043 | Keewatin Gp volcanics | 2520 | | | Keewatin | | 2840 | 2880 | 0 | 0 | | 0 | 0 |
| 22244 | 6169 | 777 | 1 | 60370.172 | Rice Lake lower unit | 7895 | | | Rice Lake | | 2840 | 2880 | 0 | 0 | | 0 | 0 |
| 22384 | 6233 | 762 | 1 | 182089.953 | Unnamed | | | | | | 2875 | 3062.5 | 0 | 0 | | 0 | 0 |
| 22396 | 6239 | 761 | 1 | 175195.859 | Unnamed | | | | | | 2875 | 3062.5 | 0 | 0 | | 0 | 0 |
| 22219 | 6157 | 781 | 1 | 19937.52 | Keewatin Unnamed 2 | 2520 | | | Keewatin | | 2880 | 2900 | 0 | 0 | | 0 | 0 |

| lith | environ | econ | measure |
|---|---|---|---|
| siliciclastic siliciclastic sedimentary ~ 0.8333\| ironstone chemical sedimentary ~ 0.1667 | fluvial indet. fluvial non-marine | | |
| sandstone siliciclastic sedimentary ~ 0.8333\|oolitic ironstone chemical sedimentary ~ 0.1667 | marine marine | | |
| sandstone siliciclastic sedimentary ~ 0.8333\|oolitic ironstone chemical sedimentary ~ 0.1667 | marine marine | | |
| sandstone siliciclastic sedimentary ~ 0.8333\|oolitic ironstone chemical sedimentary ~ 0.1667 | marine marine | | |
| ironstone chemical sedimentary ~ 0.7143\| conglomerate siliciclastic sedimentary ~ 0.1429\| sandstone siliciclastic sedimentary ~ 0.1429 | lacustrine indet. lacustrine non-marine \|floodplain fluvial non-marine \|fluvial indet. fluvial non-marine | | |
| coal organic sedimentary ~ 0.0588\| ironstone chemical sedimentary ~ 0.0588\| claystone siliciclastic sedimentary ~ 0.2941\| shale siliciclastic sedimentary ~ 0.2941\| siltstone siliciclastic sedimentary ~ 0.2941 | fluvial indet. fluvial non-marine \|delta plain fluvial non-marine | | [object Object] |
| bentonitic clay siliciclastic sedimentary ~ 0.125\| ironstone chemical sedimentary ~ 0.125\| shale siliciclastic sedimentary ~ 0.625\| ash volcanic igneous ~ 0.125 | offshore siliciclastic marine | | |
| bentonitic clay siliciclastic sedimentary ~ 0.125\| ironstone chemical sedimentary ~ 0.125\| shale siliciclastic sedimentary ~ 0.625\| ash volcanic igneous ~ 0.125 | offshore siliciclastic marine | | |
| ash volcanic igneous ~ 0.1429\| ironstone chemical sedimentary ~ 0.1429\| shale siliciclastic sedimentary ~ 0.7143 | offshore siliciclastic marine | | |
| ash volcanic igneous ~ 0.1429\| ironstone chemical sedimentary ~ 0.1429\| shale siliciclastic sedimentary ~ 0.7143 | offshore siliciclastic marine | | |
| ash volcanic igneous ~ 0.1429\| ironstone chemical sedimentary ~ 0.1429\| shale siliciclastic sedimentary ~ 0.7143 | offshore siliciclastic marine | | |
| ash volcanic igneous ~ 0.1429\| ironstone chemical sedimentary ~ 0.1429\| shale siliciclastic sedimentary ~ 0.7143 | offshore siliciclastic marine \|deep-water indet. siliciclastic marine | | |
| ash volcanic igneous ~ 0.1429\| ironstone chemical sedimentary ~ 0.1429\| shale siliciclastic sedimentary ~ 0.7143 | offshore siliciclastic marine \|deep-water indet. siliciclastic marine | | |
| shale siliciclastic sedimentary ~ 0.3333\| sandstone siliciclastic sedimentary ~ 0.0667\| limestone carbonate sedimentary ~ 0.0667\| coal organic sedimentary ~ 0.0667\| ironstone chemical sedimentary ~ 0.0667\| clay siliciclas | inferred marine marine | oil/gas indet. hydrocarbon energy \|coal coal energy | |
| shale siliciclastic sedimentary ~ 0.3571\| sandstone siliciclastic sedimentary ~ 0.0714\| limestone carbonate sedimentary ~ 0.0714\| coal organic sedimentary ~ 0.0714\| ironstone chemical sedimentary ~ 0.0714\| clay siliciclas | inferred marine marine | coal coal energy | |
| shale siliciclastic sedimentary ~ 0.3571\| sandstone siliciclastic sedimentary ~ 0.0714\| limestone carbonate sedimentary ~ 0.0714\| coal organic sedimentary ~ 0.0714\| ironstone chemical sedimentary ~ 0.0714\| clay siliciclas | inferred marine marine | coal coal energy | |
| shale siliciclastic sedimentary ~ 0.5\| limestone carbonate sedimentary ~ 0.1\| chert chemical sedimentary ~ 0.1\|red sandstone siliciclastic sedimentary ~ 0.1\|black clay siliciclastic sedimentary ~ 0.1\|shaly ironstone chemica | inferred marine marine | | |
| shale siliciclastic sedimentary ~ 0.7143\| limestone carbonate sedimentary ~ 0.1429\|nodular ironstone chemical sedimentary ~ 0.1429 | marine marine | cement construction material | [object Object] |
| shale siliciclastic sedimentary ~ 0.625\| carbonate carbonate sedimentary ~ 0.125\| sand siliciclastic sedimentary ~ 0.125\| ironstone chemical sedimentary ~ 0.125 | inferred marine marine | | |
| shale siliciclastic sedimentary ~ 0.2941\| siltstone siliciclastic sedimentary ~ 0.2941\| gypsum evaporite sedimentary ~ 0.0588\| ironstone chemical sedimentary ~ 0.0588\| quartz arenite siliciclastic sedimentary ~ 0.2941 | marine marine | | |
| shale siliciclastic sedimentary ~ 0.1429\|argillaceous limestone carbonate sedimentary ~ 0.7143\| ironstone chemical sedimentary ~ 0.1429 | inferred marine marine | | |
| shale siliciclastic sedimentary ~ 0.3125\| siltstone siliciclastic sedimentary ~ 0.3125\| sandstone siliciclastic sedimentary ~ 0.3125\| ironstone chemical sedimentary ~ 0.0625 | inferred marine marine | | |
| shale siliciclastic sedimentary ~ 0.3125\| siltstone siliciclastic sedimentary ~ 0.3125\| sandstone siliciclastic sedimentary ~ 0.3125\| ironstone chemical sedimentary ~ 0.0625 | inferred marine marine | | |
| shale siliciclastic sedimentary ~ 0.2632\| siltstone siliciclastic sedimentary ~ 0.2632\| sandstone siliciclastic sedimentary ~ 0.2632\| ironstone chemical sedimentary ~ 0.0526\|red sandstone siliciclastic sedimentary ~ 0.0526\|b | inferred marine marine | | |
| shale siliciclastic sedimentary ~ 0.2632\| siltstone siliciclastic sedimentary ~ 0.2632\| sandstone siliciclastic sedimentary ~ 0.2632\| ironstone chemical sedimentary ~ 0.0526\|red sandstone siliciclastic sedimentary ~ 0.0526\|b | inferred marine marine | | |
| siltstone siliciclastic sedimentary ~ 0.8333\| ironstone chemical sedimentary ~ 0.1667 | inferred marine marine | | |
| shale siliciclastic sedimentary ~ 0.5\| limestone carbonate sedimentary ~ 0.1\| ironstone chemical sedimentary ~ 0.1\|red sandstone siliciclastic sedimentary ~ 0.1\|black shale siliciclastic sedimentary ~ 0.1\|calcareous sandst | inferred marine marine | | |
| shale siliciclastic sedimentary ~ 0.1429\| ironstone chemical sedimentary ~ 0.1429\| limestone carbonate sedimentary ~ 0.7143 | inferred marine marine | | |
| sandstone siliciclastic sedimentary ~ 0.8333\| ironstone chemical sedimentary ~ 0.1667 | inferred marine marine | | |
| shale siliciclastic sedimentary ~ 0.1429\|red sandstone siliciclastic sedimentary ~ 0.7143\| ironstone chemical sedimentary ~ 0.1429 | inferred marine marine | | |
| red sandstone siliciclastic sedimentary ~ 0.8333\| ironstone chemical sedimentary ~ 0.1667 | inferred marine marine | | |
| red sandstone siliciclastic sedimentary ~ 0.8333\| ironstone chemical sedimentary ~ 0.1667 | inferred marine marine | | |
| conglomerate siliciclastic sedimentary ~ 0.0909\| carbonate carbonate sedimentary ~ 0.4545\| chert chemical sedimentary ~ 0.0909\| silt siliciclastic sedimentary ~ 0.0909\| ironstone chemical sedimentary ~ 0.0909\|argillaceo | inferred marine marine | | |
| shale siliciclastic sedimentary ~ 0.4545\|hematitic carbonate carbonate sedimentary ~ 0.4545\|oolitic ironstone chemical sedimentary ~ 0.0909 | inferred marine marine | | |
| shale siliciclastic sedimentary ~ 0.8333\|oolitic ironstone chemical sedimentary ~ 0.1667 | marine marine | | |
| sandstone siliciclastic sedimentary ~ 0.8333\|oolitic ironstone chemical sedimentary ~ 0.1667 | inferred marine marine | | |
| sandstone siliciclastic sedimentary ~ 0.7143\|oolitic ironstone chemical sedimentary ~ 0.1429\|glauconitic sandstone siliciclastic sedimentary ~ 0.1429 | inferred marine marine | | |
| feldspathic quartzite metasedimentary metamorphic ~ 0.5\| metaconglomerate metasedimentary metamorphic ~ 0.1\| metagraywacke metasedimentary metamorphic ~ 0.1\| ironstone chemical sedimentary ~ 0.1\| arkose silic | fluvial-lacustrine indet. fluvial non-marine \|fluvial indet. fluvial non-marine | | |
| arkose siliciclastic sedimentary ~ 0.5\| ironstone chemical sedimentary ~ 0.1\| metaconglomerate metasedimentary metamorphic ~ 0.1\| metagraywacke metasedimentary metamorphic ~ 0.1\| metavolcanic metaigneous meta | fluvial indet. fluvial non-marine \|fluvial-lacustrine indet. fluvial non-marine | | |
| conglomeratic mudstone siliciclastic sedimentary ~ 0.4167\| sandstone siliciclastic sedimentary ~ 0.0833\| diamictite siliciclastic sedimentary ~ 0.4167\| ironstone chemical sedimentary ~ 0.0833 | marine marine \|glacial indet. glacial non-marine | | [object Object] |
| mudstone siliciclastic sedimentary ~ 0.8333\| iron formation chemical sedimentary ~ 0.1667 | fluvial indet. fluvial non-marine \|inferred marine marine | | [object Object] |
| iron formation chemical sedimentary ~ 1 | inferred marine marine | | |
| amphibolite metamorphic metamorphic ~ 0.2778\| gneiss metamorphic metamorphic ~ 0.2778\| metabasalt metaigneous metamorphic ~ 0.2778\| metarhyolite metaigneous metamorphic ~ 0.0556\| iron formation chemical se | inferred marine marine | | |
| iron formation chemical sedimentary ~ 0.5\| limestone carbonate sedimentary ~ 0.5 | inferred marine marine | | |
| conglomerate siliciclastic sedimentary ~ 0.4167\| iron formation chemical sedimentary ~ 0.0833\| quartzite metasedimentary metamorphic ~ 0.0833\| shale siliciclastic sedimentary ~ 0.4167 | inferred marine marine | | |
| argillite siliciclastic sedimentary ~ 0.0909\| basalt volcanic igneous ~ 0.0909\| iron formation chemical sedimentary ~ 0.0909\| tuff volcanic igneous ~ 0.0909\|stromatolitic dolomite carbonate sedimentary ~ 0.0909\|cherty limes | inferred marine marine | | |
| shale siliciclastic sedimentary ~ 0.625\| graywacke siliciclastic sedimentary ~ 0.125\| iron formation chemical sedimentary ~ 0.125\| quartzite metasedimentary metamorphic ~ 0.125 | marine marine | | |
| iron formation chemical sedimentary ~ 0.4545\| slate metasedimentary metamorphic ~ 0.4545\|chloritic slate metasedimentary metamorphic ~ 0.0909 | inferred marine marine | | [object Object],[object Object] |
| iron formation chemical sedimentary ~ 1 | inferred marine marine | | |
| basalt volcanic igneous ~ 0.3571\| greywacke siliciclastic sedimentary ~ 0.0714\| iron formation chemical sedimentary ~ 0.0714\| rhyolite volcanic igneous ~ 0.3571\| tuff volcanic igneous ~ 0.0714\| volcaniclastic siliciclastic se | inferred marine marine | | |
| graywacke siliciclastic sedimentary ~ 0.625\| iron formation chemical sedimentary ~ 0.125\| shale siliciclastic sedimentary ~ 0.125\| quartzite metasedimentary metamorphic ~ 0.125 | marine marine | | |
| slate metasedimentary metamorphic ~ 0.4167\| iron formation chemical sedimentary ~ 0.4167\|sideritic iron formation chemical sedimentary ~ 0.0833\| chert chemical sedimentary ~ 0.0833 | inferred marine marine | | [object Object],[object Object] |
| iron formation chemical sedimentary ~ 1 | inferred marine marine | | [object Object] |
| argillite siliciclastic sedimentary ~ 0.5\| iron formation chemical sedimentary ~ 0.5 | inferred marine marine | | |
| shale siliciclastic sedimentary ~ 0.1667\| iron formation chemical sedimentary ~ 0.8333 | inferred marine marine | | |
| argillite siliciclastic sedimentary ~ 0.3846\| conglomerate siliciclastic sedimentary ~ 0.3846\| iron formation chemical sedimentary ~ 0.0769\| limestone carbonate sedimentary ~ 0.0769\| tuff volcanic igneous ~ 0.0769 | inferred marine marine | | |
| argillite siliciclastic sedimentary ~ 0.125\| dolomite carbonate sedimentary ~ 0.625\| greywacke siliciclastic sedimentary ~ 0.125\| iron formation chemical sedimentary ~ 0.125 | lacustrine indet. lacustrine non-marine | | [object Object] |
| quartz arenite siliciclastic sedimentary ~ 0.4545\| arkose siliciclastic sedimentary ~ 0.0909\| basalt volcanic igneous ~ 0.0909\| iron formation chemical sedimentary ~ 0.0909\|black mudstone siliciclastic sedimentary ~ 0.0909 | inferred marine marine | | |
| iron formation chemical sedimentary ~ 1 | inferred marine marine | | [object Object] |
| iron formation chemical sedimentary ~ 1 | inferred marine marine | | |
| iron formation chemical sedimentary ~ 0.5\| chert chemical sedimentary ~ 0.5 | inferred marine marine | | |
| basalt volcanic igneous ~ 0.1667\| iron formation chemical sedimentary ~ 0.8333 | inferred marine marine | | [object Object] |
| slate metasedimentary metamorphic ~ 0.1667\| iron formation chemical sedimentary ~ 0.8333 | inferred marine marine | | |
| iron formation chemical sedimentary ~ 1 | inferred marine marine | | |
| iron formation chemical sedimentary ~ 1 | inferred marine marine | | [object Object],[object Object] |
| iron formation chemical sedimentary ~ 0.1667\| shale siliciclastic sedimentary ~ 0.8333 | inferred marine marine | | [object Object] |
| mafic volcanic volcanic igneous ~ 0.8333\| iron formation chemical sedimentary ~ 0.1667 | inferred marine marine | | |
| slate metasedimentary metamorphic ~ 0.5556\| phosphorite chemical sedimentary ~ 0.1111\| phyllite metasedimentary metamorphic ~ 0.1111\|black slate metasedimentary metamorphic ~ 0.1111\| iron formation chemical se | inferred marine marine | | [object Object],[object Object] |
| shale siliciclastic sedimentary ~ 0.625\| iron formation chemical sedimentary ~ 0.125\| quartzite metasedimentary metamorphic ~ 0.125\|carbonaceous shale siliciclastic sedimentary ~ 0.125 | inferred marine marine | | |
| granodiorite plutonic igneous ~ 0.4545\| schist metamorphic metamorphic ~ 0.0909\| amphibolite metamorphic metamorphic ~ 0.0909\| iron formation chemical sedimentary ~ 0.0909\| metasedimentary metasedimentary met | inferred marine marine | | |
| gneiss metamorphic metamorphic ~ 0.1\| monzonite plutonic igneous ~ 0.5\| iron formation chemical sedimentary ~ 0.1\| metasedimentary metasedimentary metamorphic ~ 0.1\| metavolcanic metaigneous metamorphic ~ 0. | inferred marine marine | oil indet. hydrocarbon energy | |
| chert chemical sedimentary ~ 0.1\| dolomite carbonate sedimentary ~ 0.1\| iron formation chemical sedimentary ~ 0.1\| quartzite metasedimentary metamorphic ~ 0.1\| shale siliciclastic sedimentary ~ 0.5\| slate metasediment | inferred marine marine | | |
| iron formation chemical sedimentary ~ 1 | inferred marine marine | | [object Object] |
| metavolcanic metaigneous metamorphic ~ 0.5556\| iron formation chemical sedimentary ~ 0.1111\| metasedimentary metasedimentary metamorphic ~ 0.1111\| volcaniclastic siliciclastic sedimentary ~ 0.1111\| gabbro plutoni | inferred marine marine | | |
| arkose siliciclastic sedimentary ~ 0.2941\| greywacke siliciclastic sedimentary ~ 0.2941\| iron formation chemical sedimentary ~ 0.0588\| quartzite metasedimentary metamorphic ~ 0.0588\| shale siliciclastic sedimentary ~ 0.2 | inferred marine marine | | |
| iron formation chemical sedimentary ~ 0.8333\| chert chemical sedimentary ~ 0.1667 | inferred marine marine | | |
| graywacke siliciclastic sedimentary ~ 0.3846\| quartzite metasedimentary metamorphic ~ 0.3846\| iron formation chemical sedimentary ~ 0.0769\| marble metasedimentary metamorphic ~ 0.0769\| metavolcanic metaigneous | inferred marine marine | | |
| gneiss metamorphic metamorphic ~ 0.25\| granite plutonic igneous ~ 0.25\| migmatite metamorphic metamorphic ~ 0.25\| iron formation chemical sedimentary ~ 0.05\| marble metasedimentary metamorphic ~ 0.05\| metavolc | inferred marine marine | | |
| gneiss metamorphic metamorphic ~ 0.2381\| granite plutonic igneous ~ 0.2381\| migmatite metamorphic metamorphic ~ 0.2381\| argillite siliciclastic sedimentary ~ 0.0476\| iron formation chemical sedimentary ~ 0.0476\| mar | inferred marine marine | | |
| gneiss metamorphic metamorphic ~ 0.2381\| granite plutonic igneous ~ 0.2381\| migmatite metamorphic metamorphic ~ 0.2381\| iron formation chemical sedimentary ~ 0.0476\| marble metasedimentary metamorphic ~ 0.04 | inferred marine marine | | |
| chert chemical sedimentary ~ 0.4545\| iron formation chemical sedimentary ~ 0.4545\| metasedimentary metasedimentary metamorphic ~ 0.0909 | inferred marine marine | | |
| iron formation chemical sedimentary ~ 0.0909\| migmatite metamorphic metamorphic ~ 0.0909\| siltstone siliciclastic sedimentary ~ 0.0909\| slate metasedimentary metamorphic ~ 0.0909\| wackestone carbonate sedimentary | inferred marine marine | | |
| metavolcanic metaigneous metamorphic ~ 0.4545\| argillite siliciclastic sedimentary ~ 0.0909\|biotite gneiss metamorphic metamorphic ~ 0.0909\| iron formation chemical sedimentary ~ 0.0909\| marble metasedimentary met | inferred marine marine | | |
| metasedimentary metasedimentary metamorphic ~ 0.5556\| iron formation chemical sedimentary ~ 0.1111\| metaconglomerate metasedimentary metamorphic ~ 0.1111\| metagraywacke metasedimentary metamorphic ~ 0.1 | inferred marine marine | | |
| arkose siliciclastic sedimentary ~ 0.2632\| conglomerate siliciclastic sedimentary ~ 0.2632\| quartzite metasedimentary metamorphic ~ 0.2632\| dolomite carbonate sedimentary ~ 0.0526\| iron formation chemical sedimentary | inferred marine marine | | |
| volcaniclastic siliciclastic sedimentary ~ 0.1667\| iron formation chemical sedimentary ~ 0.8333 | inferred marine marine | | |
| arkose siliciclastic sedimentary ~ 0.2381\| conglomerate siliciclastic sedimentary ~ 0.2381\| greywacke siliciclastic sedimentary ~ 0.2381\| iron formation chemical sedimentary ~ 0.0476\| shale siliciclastic sedimentary ~ 0.238 | inferred marine marine | | |
| arkose siliciclastic sedimentary ~ 0.2381\| conglomerate siliciclastic sedimentary ~ 0.2381\| greywacke siliciclastic sedimentary ~ 0.2381\| iron formation chemical sedimentary ~ 0.0476\| shale siliciclastic sedimentary ~ 0.238 | fluvial-deltaic indet. fluvial non-marine | | [object Object] |
| chert chemical sedimentary ~ 0.1429\| iron formation chemical sedimentary ~ 0.1429\| greenstone metavolcanic metamorphic ~ 0.7143 | inferred marine marine | | [object Object] |
| arkose siliciclastic sedimentary ~ 0.2273\| conglomerate siliciclastic sedimentary ~ 0.2273\| greywacke siliciclastic sedimentary ~ 0.2273\| iron formation chemical sedimentary ~ 0.0455\| shale siliciclastic sedimentary ~ 0.227 | inferred marine marine | | |
| arkose siliciclastic sedimentary ~ 0.2941\| greywacke siliciclastic sedimentary ~ 0.2941\| iron formation chemical sedimentary ~ 0.0588\| quartzite metasedimentary metamorphic ~ 0.0588\| shale siliciclastic sedimentary ~ 0.2 | inferred marine marine | | |
| greywacke siliciclastic sedimentary ~ 0.8333\| iron formation chemical sedimentary ~ 0.1667 | inferred marine marine | | |
| iron formation chemical sedimentary ~ 1 | inferred marine marine | | |
| basalt volcanic igneous ~ 0.1667\| greywacke siliciclastic sedimentary ~ 0.1667\| iron formation chemical sedimentary ~ 0.1667\| quartzite metasedimentary metamorphic ~ 0.1667\| shale siliciclastic sedimentary ~ 0.1667\| tu | inferred marine marine | | |
| conglomerate siliciclastic sedimentary ~ 0.4545\| greywacke siliciclastic sedimentary ~ 0.4545\| iron formation chemical sedimentary ~ 0.0909 | inferred marine marine | | |
| basalt volcanic igneous ~ 0.2\| greywacke siliciclastic sedimentary ~ 0.2\| iron formation chemical sedimentary ~ 0.2\| tuff volcanic igneous ~ 0.2\| volcaniclastic siliciclastic sedimentary ~ 0.2 | inferred marine marine | | |
| basalt volcanic igneous ~ 0.25\| iron formation chemical sedimentary ~ 0.25\| tuff volcanic igneous ~ 0.25\| volcaniclastic siliciclastic sedimentary ~ 0.25 | inferred marine marine | | |
| conglomerate siliciclastic sedimentary ~ 0.3125\| greywacke siliciclastic sedimentary ~ 0.3125\| iron formation chemical sedimentary ~ 0.0625\| shale siliciclastic sedimentary ~ 0.3125 | inferred marine marine | | |
| basalt volcanic igneous ~ 0.1429\| greywacke siliciclastic sedimentary ~ 0.1429\| iron formation chemical sedimentary ~ 0.1429\| rhyolite volcanic igneous ~ 0.1429\| shale siliciclastic sedimentary ~ 0.1429\| tuff volcanic ignec | inferred marine marine | | |
| greywacke siliciclastic sedimentary ~ 0.2941\| iron formation chemical sedimentary ~ 0.0588\| rhyolite volcanic igneous ~ 0.0588\| shale siliciclastic sedimentary ~ 0.2941\| volcaniclastic siliciclastic sedimentary ~ 0.2941 | inferred marine marine | | |
| basalt volcanic igneous ~ 0.1667\| greywacke siliciclastic sedimentary ~ 0.1667\| iron formation chemical sedimentary ~ 0.1667\| rhyolite volcanic igneous ~ 0.1667\| tuff volcanic igneous ~ 0.1667\| volcaniclastic siliciclastic se | inferred marine marine | | |
| basalt volcanic igneous ~ 0.2\| iron formation chemical sedimentary ~ 0.2\| rhyolite volcanic igneous ~ 0.2\| tuff volcanic igneous ~ 0.2\| volcaniclastic siliciclastic sedimentary ~ 0.2 | inferred marine marine | | |
| basalt volcanic igneous ~ 0.1\| greywacke siliciclastic sedimentary ~ 0.1\| iron formation chemical sedimentary ~ 0.1\| quartzite metasedimentary metamorphic ~ 0.5\| rhyolite volcanic igneous ~ 0.1\| shale siliciclastic sedimen | inferred marine marine | | |
| basalt volcanic igneous ~ 0.25\| iron formation chemical sedimentary ~ 0.25\| tuff volcanic igneous ~ 0.25\| volcaniclastic siliciclastic sedimentary ~ 0.25 | inferred marine marine | | |
| basalt volcanic igneous ~ 0.2\| iron formation chemical sedimentary ~ 0.2\| rhyolite volcanic igneous ~ 0.2\| tuff volcanic igneous ~ 0.2\| volcaniclastic siliciclastic sedimentary ~ 0.2 | inferred marine marine | | |
| tuff volcanic igneous ~ 0.0909\| basalt volcanic igneous ~ 0.4545\| iron formation chemical sedimentary ~ 0.4545 | inferred marine marine | | |
| iron formation chemical sedimentary ~ 0.1111\| rhyolite volcanic igneous ~ 0.1111\| tuff volcanic igneous ~ 0.1111\| volcaniclastic siliciclastic sedimentary ~ 0.1111\| basalt volcanic igneous ~ 0.5556 | inferred marine marine | | |
| greywacke siliciclastic sedimentary ~ 0.4167\| iron formation chemical sedimentary ~ 0.0833\| tuff volcanic igneous ~ 0.0833\| volcaniclastic siliciclastic sedimentary ~ 0.4167 | inferred marine marine | | |

| notes | color | text_color | t_int_id | t_int_name | t_int_age | t_prop | units_above | b_int_id | b_int_name | b_int_age | b_prop | units_below | strat_name_long |
|---|---|---|---|---|---|---|---|---|---|---|---|---|---|
| High-level Fluvial Deposits | #A57B54 | #000000 | 4 | Pleistocene | 0.0117 | 0.25001 | 0 | 12 | Zanclean | 5.333 | 0 | 0 | |
| S.S. & Envr: https://era.library.ualberta.ca/items/bb399a54-2fb3-42f4-a95a-2a4252a62212 | #FCF768 | #000000 | 37 | Santonian | 83.6 | 0.75 | 37329 | 37 | Santonian | 86.3 | 0 | 37327 | Badheart Formation |
| S.S. & Envr: https://era.library.ualberta.ca/items/bb399a54-2fb3-42f4-a95a-2a4252a62212 | #FCF768 | #000000 | 37 | Santonian | 83.6 | 0.66667 | 37147 | 37 | Santonian | 86.3 | 0 | 37145 | Badheart Formation |
| S.S. & Envr: https://era.library.ualberta.ca/items/bb399a54-2fb3-42f4-a95a-2a4252a62212 | #FCF768 | #000000 | 37 | Santonian | 83.6 | 0.25 | 37279 | 37 | Santonian | 86.3 | 0 | 37277 | Badheart Formation |
| environ- MNGS | #FCF768 | #000000 | 40 | Cenomanian | 93.9 | 1 | 0 | 40 | Cenomanian | 100.5 | 0.75 | 0 | Windrow Formation |
| environ- Flores et al 2005 | #FCF768 | #000000 | 40 | Cenomanian | 93.9 | 0.83333 | 0 | 40 | Cenomanian | 100.5 | 0.66667 | 5238 | Corwin Formation |
| environ- Shurr & Ridgley 2002 | #999999 | #000000 | 40 | Cenomanian | 93.9 | 0.75 | 9496 | 40 | Cenomanian | 100.5 | 0 | 9499 | Belle Fourche Formation |
| environ- Caritat et al 1997 | #999999 | #000000 | 40 | Cenomanian | 93.9 | 0.75 | 9595 | 40 | Cenomanian | 100.5 | 0 | 9598 | Belle Fourche Formation |
| environ- Shurr & Ridgley 2002 | #999999 | #000000 | 40 | Cenomanian | 93.9 | 0.75 | 9626 | 40 | Cenomanian | 100.5 | 0 | 9629 | Belle Fourche Formation |
| environ- Caritat et al 1997 | #999999 | #000000 | 40 | Cenomanian | 93.9 | 0.75 | 9712 | 40 | Cenomanian | 100.5 | 0 | 9715 | Belle Fourche Formation |
| environ- Shurr & Ridgley 2002 | #999999 | #000000 | 40 | Cenomanian | 93.9 | 0.75 | 9794 | 40 | Cenomanian | 100.5 | 0 | 9796 | Belle Fourche Formation |
| environ- Skolnick 1958 (Mowry) and Shurr & Ridgley 2002 (Belle Fourche) | #999999 | #000000 | 40 | Cenomanian | 93.9 | 0.75 | 9856 | 42 | Albian | 113 | 0.83333 | 9858 | Belle Fourche Formation |
| environ- Skolnick 1958 (Mowry) and Shurr & Ridgley 2002 (Belle Fourche) | #999999 | #000000 | 40 | Cenomanian | 93.9 | 0.75 | 9901 | 42 | Albian | 113 | 0.83333 | 9903 | Belle Fourche Formation |
| | #999999 | #000000 | 139 | Desmoinesian | 306 | 0.33333 | 0,11397 | 141 | Morrowan | 323.2 | 0.25 | 0,11397 | Saginaw Formation |
| | #999999 | #000000 | 139 | Desmoinesian | 306 | 0.33333 | 0,11420 | 141 | Morrowan | 323.2 | 0.25 | 0,11420 | Saginaw Formation |
| | #999999 | #000000 | 139 | Desmoinesian | 306 | 0.33333 | 0,11487 | 141 | Morrowan | 323.2 | 0.25 | 0,11487 | Saginaw Formation |
| | #999999 | #000000 | 140 | Atokan | 312 | 1 | 15283 | 140 | Atokan | 319 | 0.75 | 15285 | Atoka Formation |
| | #999999 | #000000 | 144 | Osagean | 343.5 | 0.25 | 12235 | 144 | Osagean | 351.5 | 0.125 | 12223 | New Providence Shale |
| | #999999 | #000000 | 99 | Givetian | 382.7 | 0.83332 | 0 | 99 | Givetian | 387.7 | 0.66666 | 0 | Olentangy Shale |
| S.S. Desc: http://archives.datapages.com/data/cspg/data/024/024004/0485.htm | #FCF768 | #000000 | 100 | Eifelian | 387.7 | 0.66668 | 34908 | 102 | Emsian | 407.6 | 0.75 | 34906 | Bird Fiord Formation |
| | #4D52E7 | #E0E0E0 | 100 | Eifelian | 387.7 | 0.65 | 8234 | 100 | Eifelian | 393.3 | 0 | 8237 | Buttermilk Falls Limestone |
| | #A57B54 | #000000 | 108 | Ludlow | 423 | 0.83333 | 271 | 236 | Rhuddanian | 443.8 | 0 | 276 | Rockwood Formation |
| | #A57B54 | #000000 | 108 | Ludlow | 423 | 0.83334 | 430 | 236 | Rhuddanian | 443.8 | 0 | 435 | Rockwood Formation |
| | #999999 | #000000 | 108 | Ludlow | 423 | 0.66667 | 272 | 234 | Telychian | 438.5 | 0 | 275 | Clinton Formation |
| | #999999 | #000000 | 108 | Ludlow | 423 | 0.66666 | 431 | 234 | Telychian | 438.5 | 0 | 434 | Clinton Formation |
| | #999999 | #000000 | 109 | Wenlock | 427.4 | 0.75 | 1433 | 109 | Wenlock | 433.4 | 0 | 1436 | Mifflintown Formation |
| | #A57B54 | #000000 | 109 | Wenlock | 427.4 | 0.25 | 0 | 234 | Telychian | 438.5 | 0.75 | 0 | Clinton Formation |
| | #4D52E7 | #E0E0E0 | 234 | Telychian | 433.4 | 1 | 12392 | 234 | Telychian | 438.5 | 0.75 | 0 | Dayton Formation |
| | #FCF768 | #000000 | 234 | Telychian | 433.4 | 0.33333 | 1992 | 234 | Telychian | 438.5 | 0.16667 | 1992 | Cresaptown Sandstone Member |
| S.S. Desc: https://pubs.geoscienceworld.org/sepm/jsedres/article/30/1/1/95454/Petrography-and-origin-of-the-Tuscarora-Rose-Hill | #FCF768 | #000000 | 234 | Telychian | 433.4 | 0.25 | 1221 | 234 | Telychian | 438.5 | 0.16667 | 1221 | Cacapon Sandstone Member |
| S.S. Desc: https://pubs.geoscienceworld.org/sepm/jsedres/article/30/1/1/95454/Petrography-and-origin-of-the-Tuscarora-Rose-Hill | #FCF768 | #000000 | 234 | Telychian | 433.4 | 0.25 | 1321 | 234 | Telychian | 438.5 | 0.16667 | 1321 | Cacapon Sandstone Member |
| S.S. Desc: https://pubs.geoscienceworld.org/sepm/jsedres/article/30/1/1/95454/Petrography-and-origin-of-the-Tuscarora-Rose-Hill | #FCF768 | #000000 | 234 | Telychian | 433.4 | 0.25 | 1711 | 234 | Telychian | 438.5 | 0.16667 | 1711 | Cacapon Sandstone Member |
| | #4D52E7 | #E0E0E0 | 235 | Aeronian | 438.5 | 1 | 0 | 235 | Aeronian | 440.8 | 0 | 0 | Brassfield Formation |
| | #999999 | #000000 | 160 | Richmondian | 445.2 | 0.83334 | 0 | 160 | Richmondian | 449 | 0.66666 | 11217 | Neda Formation |
| | #999999 | #000000 | 166 | Blackriverian | 456.5 | 0.25 | 2964 | 166 | Blackriverian | 458.4 | 0 | 2966 | Glenwood Member |
| | #FCF768 | #000000 | 127 | Dresbachian | 496.8 | 0.75 | 2959 | 127 | Dresbachian | 501 | 0 | 0 | Mount Simon Sandstone |
| | #FCF768 | #000000 | 127 | Dresbachian | 496.8 | 0.75 | 3005 | 127 | Dresbachian | 501 | 0 | 0 | Mount Simon Sandstone |
| limestone is rare | #FCF768 | #000000 | 263 | Ediacaran | 538.8 | 0.3 | 0 | 263 | Ediacaran | 635 | 0.1 | 0 | Chestnut Hill Formation |
| | #FCF768 | #000000 | 263 | Ediacaran | 538.8 | 0.3 | 0 | 263 | Ediacaran | 635 | 0.1 | 0 | Chestnut Hill Formation |
| | #A57B54 | #000000 | 264 | Cryogenian | 635 | 1 | 0 | 264 | Cryogenian | 720 | 0 | 13874 | Kingston Peak Formation |
| Rapitan formation | #F58D2C | #000000 | 264 | Cryogenian | 635 | 0.75 | 0 | 264 | Cryogenian | 720 | 0.66667 | 0 | Sayunei Formation |
| | #777777 | #FFFFFF | 201 | Aphebian | 1600 | 0.9 | 0 | 201 | Aphebian | 2500 | 0.8 | 22044 | Wabush Lake Formation |
| Moppin and Gold Hill complexes, Pecos complex; thickness mostly unconstrained; arc complexes | #004703 | #000000 | 269 | Statherian | 1600 | 0.38 | 0 | 269 | Statherian | 1800 | 0.25 | 0 | |
| Belcher Group | #777777 | #FFFFFF | 201 | Aphebian | 1600 | 0.84 | 22289 | 201 | Aphebian | 2500 | 0.83 | 22291 | Kipalu Formation |
| Mistassini Group | #FCF768 | #000000 | 201 | Aphebian | 1600 | 0.83333 | 0 | 201 | Aphebian | 2500 | 0.66667 | 0 | Temiscamie Formation |
| The age of the IF is constrained between 1870 Ma, from U-Pb in baddeleyite from the Flaherty basalt correlative with the basalt overlying the Nastapoka IF (Hamilton et al., 2009), and 2025Â±25 Ma, from U-Pb in uraniferous | #34A038 | #000000 | 201 | Aphebian | 1600 | 0.78 | 0 | 201 | Aphebian | 2500 | 0.7 | 22304 | Nastapoka Group |
| | #999999 | #000000 | 270 | Orosirian | 1800 | 1 | 0 | 270 | Orosirian | 2050 | 0.8 | 43002 | Animikie Group |
| | #A57B54 | #000000 | 259 | Paleoproterozoic | 1600 | 0.755 | 33359 | 259 | Paleoproterozoic | 2500 | 0.7325 | 33357 | Stambaugh Formation |
| | #F58D2C | #000000 | 270 | Orosirian | 1800 | 0.8 | 5942 | 270 | Orosirian | 2050 | 0.75 | 5942 | Emily Member |
| | #34A038 | #000000 | 270 | Orosirian | 1800 | 0.8 | 22345 | 270 | Orosirian | 2050 | 0.68 | 0 | Wasekwan Group |
| | #FCF768 | #000000 | 270 | Orosirian | 1800 | 0.8 | 43003 | 270 | Orosirian | 2050 | 0.5 | 43001 | North Range Group |
| | #999999 | #000000 | 259 | Paleoproterozoic | 1600 | 0.72 | 33357 | 259 | Paleoproterozoic | 2500 | 0.7 | 33356 | Riverton Iron Formation |
| | #F58D2C | #000000 | 270 | Orosirian | 1800 | 0.75 | 5942 | 270 | Orosirian | 2050 | 0.7 | 5945 | Trommald Formation |
| | #F58D2C | #000000 | 270 | Orosirian | 1800 | 0.75 | 5966 | 270 | Orosirian | 2050 | 0.625 | 5966 | Virginia Formation |
| | #F58D2C | #000000 | 270 | Orosirian | 1800 | 0.75 | 5971 | 270 | Orosirian | 2050 | 0.69 | 0 | Gunflint Iron Formation |
| | #999999 | #000000 | 270 | Orosirian | 1800 | 0.75 | 22579 | 270 | Orosirian | 2050 | 0.69 | 22577 | Gunflint Iron Formation |
| Hurwitz Group | #4D52E7 | #E0E0E0 | 201 | Aphebian | 1600 | 0.7 | 22389 | 201 | Aphebian | 2500 | 0.6 | 22387,22391 | Hurwitz Group |
| age and thickness updated from Rainbird et al. 2010 | #FCF768 | #000000 | 270 | Orosirian | 1800 | 0.69 | 0 | 271 | Rhyacian | 2300 | 0.48 | 0 | Ketyet River Group |
| | #F58D2C | #000000 | 259 | Paleoproterozoic | 1600 | 0.6875 | 6168,6169 | 259 | Paleoproterozoic | 2500 | 0.625 | 6172 | Ironwood Iron Formation |
| | #F58D2C | #000000 | 270 | Orosirian | 1800 | 0.5 | 5987 | 270 | Orosirian | 2050 | 0.4 | 5989 | Trommald Formation |
| | #F58D2C | #000000 | 270 | Orosirian | 1800 | 0.46875 | 5966 | 270 | Orosirian | 2050 | 0.40625 | 5968 | Biwabik Iron Formation |
| Knob Lake Group of Kaniapiskau Supergroup | #777777 | #FFFFFF | 201 | Aphebian | 1600 | 0.62333 | 22317 | 201 | Aphebian | 2500 | 0.58667 | 22319 | Sokoman Formation |
| | #F58D2C | #000000 | 259 | Paleoproterozoic | 1600 | 0.6 | 0 | 259 | Paleoproterozoic | 2500 | 0.55 | 0 | |
| | #F58D2C | #000000 | 259 | Paleoproterozoic | 1600 | 0.6 | 0 | 259 | Paleoproterozoic | 2500 | 0.55 | 6203 | Amasa Formation |
| | #F58D2C | #000000 | 259 | Paleoproterozoic | 1600 | 0.6 | 0 | 259 | Paleoproterozoic | 2500 | 0.55 | 6238 | Vulcan Iron Formation |
| Knob Lake Group of Kaniapiskau Supergroup | #777777 | #FFFFFF | 201 | Aphebian | 1600 | 0.58667 | 22318 | 201 | Aphebian | 2500 | 0.55 | 22320 | Ruth Formation |
| | #438A47 | #000000 | 270 | Orosirian | 1800 | 0.25 | 5948 | 270 | Orosirian | 2050 | 0.125 | 0 | Randall Formation |
| | #999999 | #000000 | 259 | Paleoproterozoic | 1600 | 0.56 | 0 | 259 | Paleoproterozoic | 2500 | 0.53 | 6212 | Michigamme Formation |
| foreland basin deposit; thickness estimated | #999999 | #000000 | 270 | Orosirian | 1800 | 0.2 | 42715 | 271 | Rhyacian | 2300 | 0.85 | 42713 | |
| arc-related rocks; thickness general | #5A4633 | #000000 | 259 | Paleoproterozoic | 1600 | 0.5 | 0 | 259 | Paleoproterozoic | 2500 | 0 | 0 | |
| altered monzonite, metasediments of trans-hudson orogen | #5A4633 | #000000 | 259 | Paleoproterozoic | 1600 | 0.5 | 0 | 259 | Paleoproterozoic | 2500 | 0 | 0 | |
| Povungnituk Group | #777777 | #FFFFFF | 201 | Aphebian | 1600 | 0.495 | 22330 | 201 | Aphebian | 2500 | 0.4125 | 0 | Povungnituk Group |
| | #F58D2C | #000000 | 259 | Paleoproterozoic | 1600 | 0.47 | 6212 | 259 | Paleoproterozoic | 2500 | 0.45 | 6215,6216 | Negaunee Iron Formation |
| Trans-Hudson Orogen arc rocks; thickness estimated | #438A47 | #000000 | 259 | Paleoproterozoic | 1600 | 0.4 | 0 | 259 | Paleoproterozoic | 2500 | 0 | 0 | |
| Assean Group | #F58D2C | #000000 | 201 | Aphebian | 1600 | 0.375 | 0 | 201 | Aphebian | 2500 | 0.25 | 22340 | |
| | #E93B3B | #000000 | 272 | Siderian | 2300 | 0.08 | 41904 | 272 | Siderian | 2500 | 0.05 | 41902 | Benchmark Iron Formation |
| pre-Yellowknife sedimentary and volcanic rocks | #5A4633 | #000000 | 260 | Neoarchean | 2500 | 1 | 0 | 260 | Neoarchean | 2800 | 0.6 | 0 | |
| metasediments and volcanics subordinate to igneous rocks | #004703 | #000000 | 260 | Neoarchean | 2500 | 0.88 | 0 | 261 | Mesoarchean | 3200 | 0.25 | 0 | |
| sediments subsidiary to plutonic rocks, difficult to separate chronostratigraphically | #004703 | #000000 | 260 | Neoarchean | 2500 | 0.86 | 0 | 261 | Mesoarchean | 3200 | 0.27 | 0 | |
| metavolcanics and metasediments subsidiary to igneous; difficult to separate chronostratigraphically here | #004703 | #000000 | 260 | Neoarchean | 2500 | 0.85 | 0 | 261 | Mesoarchean | 3200 | 0.25 | 0 | |
| | #5E3006 | #000000 | 260 | Neoarchean | 2500 | 0.75 | 0 | 260 | Neoarchean | 2800 | 0.6 | 0 | Matlock Iron Formation |
| | #FCF768 | #000000 | 260 | Neoarchean | 2500 | 0.75 | 18213 | 260 | Neoarchean | 2800 | 0.25 | 0 | |
| Minor relative to granitic rocks | #5A4633 | #000000 | 260 | Neoarchean | 2500 | 0.66667 | 0 | 261 | Mesoarchean | 3200 | 0.5 | 0 | |
| | #FCF768 | #000000 | 260 | Neoarchean | 2500 | 0.65 | 0 | 260 | Neoarchean | 2800 | 0.52 | 0 | |
| | #FCF768 | #000000 | 260 | Neoarchean | 2500 | 0.5 | 21983 | 260 | Neoarchean | 2800 | 0.4 | 21984 | Wilson Island Group |
| | #F58D2C | #000000 | 260 | Neoarchean | 2500 | 0.45 | 5960 | 260 | Neoarchean | 2800 | 0.4 | 5963 | Soudan Iron Formation Member |
| | #F58D2C | #000000 | 260 | Neoarchean | 2500 | 0.45 | 0 | 260 | Neoarchean | 2800 | 0.2 | 22574 | |
| Timiskaming Group | #F58D2C | #000000 | 260 | Neoarchean | 2500 | 0.4333 | 22174 | 260 | Neoarchean | 2800 | 0.4111 | 0 | Timiskaming Group |
| | #438A47 | #000000 | 260 | Neoarchean | 2500 | 0.4 | 5962 | 260 | Neoarchean | 2800 | 0.2 | 0 | Ely Greenstone |
| | #F58D2C | #000000 | 260 | Neoarchean | 2500 | 0.4 | 0 | 260 | Neoarchean | 2800 | 0.2 | 22231 | |
| Oxford Group | #F58D2C | #000000 | 260 | Neoarchean | 2500 | 0.4 | 0 | 260 | Neoarchean | 2800 | 0.3 | 22260 | Oxford Lake Group |
| | #F58D2C | #000000 | 260 | Neoarchean | 2500 | 0.35 | 0 | 260 | Neoarchean | 2800 | 0.2 | 22097 | |
| Steeprick Group | #F58D2C | #000000 | 260 | Neoarchean | 2500 | 0.35 | 22209 | 260 | Neoarchean | 2800 | 0.3 | 22211 | Jolliffe Ore Zone Formation |
| may be Aphebian | #634C2E | #000000 | 260 | Neoarchean | 2500 | 0.33333 | 0 | 260 | Neoarchean | 2800 | 0.16667 | 0 | |
| | #F58D2C | #000000 | 260 | Neoarchean | 2500 | 0.3 | 0 | 260 | Neoarchean | 2800 | 0.25 | 22136 | |
| | #34A038 | #000000 | 260 | Neoarchean | 2500 | 0.3 | 22226 | 261 | Mesoarchean | 3200 | 0.8 | 22228 | Keewatin Group |
| | #34A038 | #000000 | 260 | Neoarchean | 2500 | 0.2 | 22212,22213 | 261 | Mesoarchean | 3200 | 0.8 | 22215 | Keewatin Group |
| | #F58D2C | #000000 | 260 | Neoarchean | 2500 | 0.2 | 22235 | 261 | Mesoarchean | 3200 | 0.9 | 22237 | |
| Keewatin Group | #34A038 | #000000 | 260 | Neoarchean | 2500 | 0.2 | 0 | 261 | Mesoarchean | 3200 | 0.8 | 0 | Keewatin Group |
| Keewatin Group | #F58D2C | #000000 | 260 | Neoarchean | 2500 | 0.2 | 0 | 261 | Mesoarchean | 3200 | 0.9 | 22185 | Keewatin Group |
| | #34A038 | #000000 | 260 | Neoarchean | 2500 | 0.2 | 22230 | 261 | Mesoarchean | 3200 | 0.8 | 0 | Keewatin Group |
| | #438A47 | #000000 | 260 | Neoarchean | 2500 | 0.2 | 22575 | 261 | Mesoarchean | 3200 | 0.8 | 22573 | Keewatin Group |
| | #634C2E | #000000 | 132 | Archean | 2500 | 0.83333 | 0 | 132 | Archean | 4000 | 0.66667 | 0 | |
| Keewatin Group | #34A038 | #000000 | 261 | Mesoarchean | 2800 | 0.9 | 22183 | 261 | Mesoarchean | 3200 | 0.8 | 0 | Keewatin Group |
| | #34A038 | #000000 | 261 | Mesoarchean | 2800 | 0.9 | 22243 | 261 | Mesoarchean | 3200 | 0.8 | 0 | Rice Lake Group |
| | #34A038 | #000000 | 132 | Archean | 2500 | 0.75 | 22383 | 132 | Archean | 4000 | 0.625 | 0 | |
| | #34A038 | #000000 | 132 | Archean | 2500 | 0.75 | 22395 | 132 | Archean | 4000 | 0.625 | 0 | |
| Keewatin Group | #F58D2C | #000000 | 261 | Mesoarchean | 2800 | 0.8 | 22218 | 261 | Mesoarchean | 3200 | 0.75 | 22220 | Keewatin Group |

| refs | clat | clng | t_plat | t_plng | b_plat | b_plng |
|---|---|---|---|---|---|---|
| 1 | 34.963 | -87.379 | 35.328 | -86.595 | 35.941 | -85.206 |
| 12 | 54.826 | -115.211 | 64.052 | -66.867 | 63.639 | -67.164 |
| 12 | 55.88 | -120.939 | 66.214 | -73.122 | 65.773 | -73.43 |
| 12 | 54.254 | -120.18 | 64.262 | -73.69 | 64.096 | -73.757 |
| 1 | 42.751 | -92.798 | 47.422 | -48.951 | 47.426 | -48.64 |
| 1 | 69.847 | -159.802 | 79.742 | -133.305 | 79.306 | -132.818 |
| 1 | 47.279 | -103.421 | 53.457 | -58.644 | 53.218 | -57.784 |
| 1 | 48.383 | -102.95 | 54.467 | -57.715 | 54.255 | -56.95 |
| 1 | 47.246 | -100.575 | 53.008 | -55.505 | 52.866 | -54.624 |
| 1 | 48.849 | -100.8 | 54.604 | -55.148 | 54.471 | -54.409 |
| 1 | 46.266 | -102.95 | 52.399 | -58.474 | 52.166 | -57.526 |
| 1\|1 | 44.95 | -102.878 | 51.1 | -58.831 | 51.032 | -56.109 |
| 1\|1 | 45.4 | -100.34 | 51.171 | -55.893 | 51.176 | -53.18 |
| 1 | 43.825 | -84.963 | -2.628 | -19.215 | -14.674 | -28.617 |
| 1 | 42.421 | -85.683 | -3.684 | -20.281 | -15.575 | -29.858 |
| 1 | 42.416 | -83.803 | -4.26 | -19.016 | -16.317 | -28.638 |
| 1 | 39.641 | -103.333 | 0.564 | -33.305 | -0.822 | -34.46 |
| 1 | 39.091 | -85.333 | -32.021 | -50.351 | -32.272 | -50.692 |
| 1 | 39.433 | -84.116 | -32.878 | -51.682 | -32.985 | -51.843 |
| 11 | 76.759 | -86.262 | 1.249 | -42.839 | 1.105 | -48.835 |
| 1 | 40.961 | -75.81 | -33.277 | -45.442 | -33.283 | -45.704 |
| 1 | 36.45 | -83.575 | -32.291 | -75.418 | -30.469 | -109.01 |
| 1 | 36.333 | -83.066 | -32.445 | -74.948 | -30.722 | -108.611 |
| 1 | 36.45 | -83.575 | -32.18 | -76.527 | -33.838 | -103.11 |
| 1 | 36.333 | -83.066 | -32.341 | -76.06 | -34.072 | -102.681 |
| 1 | 38.092 | -79.772 | -32.506 | -80.585 | -33.413 | -89.189 |
| 1 | 43.116 | -75.778 | -28.482 | -82.279 | -28.861 | -87.408 |
| 1 | 41.041 | -83.613 | -29.979 | -91.986 | -29.986 | -94.376 |
| 1 | 39.04 | -79.154 | -32.51 | -95.018 | -32.398 | -96.653 |
| 1 | 38 | -81.611 | -33.055 | -98.337 | -32.979 | -99.157 |
| 1 | 38.445 | -80.986 | -32.738 | -97.643 | -32.666 | -98.461 |
| 1 | 39.141 | -80.075 | -32.216 | -96.641 | -32.15 | -97.455 |
| 1 | 36.03 | -86.753 | -33.379 | -106.18 | -31.891 | -109.154 |
| 1 | 43.733 | -87.845 | -20.718 | -111.476 | -20.132 | -112.093 |
| 1 | 42.738 | -95.85 | -4.688 | -116.121 | -4.078 | -115.863 |
| 1 | 42.738 | -95.85 | 21.912 | -113.125 | 22.081 | -115.39 |
| 1 | 42.511 | -94.241 | 20.795 | -112.638 | 20.966 | -114.899 |
| 1 | 40.686 | -74.875 | 0 | 0 | 0 | 0 |
| 1 | 40.993 | -74.537 | 0 | 0 | 0 | 0 |
| 1 | 36.453 | -116.92 | 0 | 0 | 0 | 0 |
| 10 | 62.81 | -127.445 | 0 | 0 | 0 | 0 |
| 2 | 52.919 | -66.869 | 0 | 0 | 0 | 0 |
| 1 | 35.463 | -106.076 | -27.436 | -176.991 | 0 | 0 |
| 2 | 56.416 | -79.251 | 0 | 0 | 0 | 0 |
| 2 | 50.065 | -74.366 | 0 | 0 | 0 | 0 |
| 2 | 56.247 | -76.352 | 0 | 0 | 0 | 0 |
| 1 | 45.616 | -94.377 | 0 | 0 | 0 | 0 |
| 1 | 46.034 | -87.813 | 0 | 0 | 0 | 0 |
| 1 | 46.633 | -94.091 | 0 | 0 | 0 | 0 |
| 2 | 56.534 | -100.906 | 0 | 0 | 0 | 0 |
| 1 | 45.616 | -94.377 | 0 | 0 | 0 | 0 |
| 1 | 46.034 | -87.813 | 0 | 0 | 0 | 0 |
| 1 | 46.633 | -94.091 | 0 | 0 | 0 | 0 |
| 1 | 47.883 | -91.716 | 0 | 0 | 0 | 0 |
| 1 | 47.845 | -90.428 | 0 | 0 | 0 | 0 |
| 2 | 48.551 | -89.204 | 0 | 0 | 0 | 0 |
| 2 | 61.536 | -97.798 | 0 | 0 | 0 | 0 |
| 2 | 64.062 | -84.821 | 0 | 0 | 0 | 0 |
| 1 | 46.366 | -89.682 | 0 | 0 | 0 | 0 |
| 1 | 46.415 | -92.678 | 0 | 0 | 0 | 0 |
| 1 | 47.883 | -91.716 | 0 | 0 | 0 | 0 |
| 2 | 54.947 | -67.006 | 0 | 0 | 0 | 0 |
| 1 | 46.207 | -89.039 | 0 | 0 | 0 | 0 |
| 1 | 46.347 | -88.191 | 0 | 0 | 0 | 0 |
| 1 | 46.034 | -87.813 | 0 | 0 | 0 | 0 |
| 2 | 54.947 | -67.006 | 0 | 0 | 0 | 0 |
| 1 | 46.633 | -94.091 | 0 | 0 | 0 | 0 |
| 1 | 46.578 | -87.64 | 0 | 0 | 0 | 0 |
| 1 | 45.435 | -112.118 | 0 | 0 | 0 | 0 |
| 1 | 47.279 | -103.421 | 0 | 0 | 0 | 0 |
| 1 | 48.383 | -102.95 | 0 | 0 | 0 | 0 |
| 2 | 61.144 | -74.836 | 0 | 0 | 0 | 0 |
| 1 | 46.578 | -87.64 | 0 | 0 | 0 | 0 |
| 1 | 46.266 | -102.95 | 0 | 0 | 0 | 0 |
| 2 | 55.746 | -97.893 | 0 | 0 | 0 | 0 |
| 1 | 43.949 | -104.206 | 0 | 0 | 0 | 0 |
| 12 | 60.756 | -115.398 | 0 | 0 | 0 | 0 |
| 1 | 43.6 | -109.836 | -18.666 | -168.846 | 0 | 0 |
| 1 | 43.579 | -108.382 | -23.284 | -169.024 | 0 | 0 |
| 1 | 42.986 | -108.398 | -23.46 | -169.641 | 0 | 0 |
| 1 | 43.267 | -96.152 | 0 | 0 | 0 | 0 |
| 2 | 55.168 | -87.152 | 0 | 0 | 0 | 0 |
| 1 | 41.65 | -106.911 | 0 | 0 | 0 | 0 |
| 1 | 43.949 | -104.206 | 0 | 0 | 0 | 0 |
| 2 | 62.904 | -109.367 | 0 | 0 | 0 | 0 |
| 1 | 47.883 | -91.716 | 0 | 0 | 0 | 0 |
| 2 | 48.551 | -89.204 | 0 | 0 | 0 | 0 |
| 2 | 48.151 | -80.029 | 0 | 0 | 0 | 0 |
| 1 | 47.883 | -91.716 | 0 | 0 | 0 | 0 |
| 2 | 49.14 | -94.333 | 0 | 0 | 0 | 0 |
| 2 | 54.611 | -97.762 | 0 | 0 | 0 | 0 |
| 2 | 46.384 | -82.652 | 0 | 0 | 0 | 0 |
| 2 | 49.252 | -87.915 | 0 | 0 | 0 | 0 |
| 2 | 71.811 | -79.157 | 0 | 0 | 0 | 0 |
| 2 | 52.233 | -78.499 | 0 | 0 | 0 | 0 |
| 2 | 48.929 | -93.863 | 0 | 0 | 0 | 0 |
| 2 | 49.252 | -87.915 | 0 | 0 | 0 | 0 |
| 2 | 51.124 | -92.517 | 0 | 0 | 0 | 0 |
| 2 | 48.151 | -80.029 | 0 | 0 | 0 | 0 |
| 2 | 48.603 | -81.279 | 0 | 0 | 0 | 0 |
| 2 | 49.14 | -94.333 | 0 | 0 | 0 | 0 |
| 2 | 48.551 | -89.204 | 0 | 0 | 0 | 0 |
| 2 | 65.892 | -90.565 | 0 | 0 | 0 | 0 |
| 2 | 48.603 | -81.279 | 0 | 0 | 0 | 0 |
| 2 | 50.338 | -95.991 | 0 | 0 | 0 | 0 |
| 2 | 60.468 | -103.182 | 0 | 0 | 0 | 0 |
| 2 | 61.536 | -97.798 | 0 | 0 | 0 | 0 |
| 2 | 48.803 | -92.775 | 0 | 0 | 0 | 0 |
