## Supplemental Table 2 for "Did Iron Suppress Eukaryote Emergence and Early Radiation?"

SI_CSV_File_2_Ferruginous_keyword_units

| unit_id | section_id | col_id | project_id | col_area | unit_name | strat_name_id | Mbr | Fm | Gp | SGp | t_age | b_age | max_thick | min_thick | outcrop | pbdb_collections | pbdb_occurrences |
|---|---|---|---|---|---|---|---|---|---|---|---|---|---|---|---|---|---|
| **4913** | 1095 | 130 | 1 | 5799.331 | Cape May Fm | 311 | | Cape May | | | 0.0117 | 0.5896 | 15 | 1 | surface | 4 | 8 |
| **4612** | 899 | 114 | 1 | 4579.002 | Cape May Fm | 311 | | Cape May | | | 0.4398 | 0.8678 | 15 | 1 | both | 0 | 0 |
| **4651** | 910 | 115 | 1 | 4926.812 | Cape May Fm | 311 | | Cape May | | | 0.4398 | 0.8678 | 15 | 1 | surface | 0 | 0 |
| **4955** | 1112 | 132 | 1 | 6998.652 | Cape May Fm | 311 | | Cape May | | | 0.4398 | 0.8678 | 15 | 1 | surface | 3 | 5 |
| **4616** | 903 | 114 | 1 | 4579.002 | Beacon Hill Gravel | 4868 | | Beacon Hill Gravel | | | 2.58 | 3.345 | 12 | 0 | surface | 0 | 0 |
| **6695** | 1666 | 232 | 1 | 5425.838 | Pullen Fm | 1662 | | Pullen | Wildcat | | 3.43 | 5.333 | 244 | 0 | surface | 0 | 0 |
| **6706** | 1668 | 233 | 1 | 6428.512 | Pullen Fm | 1662 | | Pullen | Wildcat | | 3.43 | 5.8113 | 345 | 0 | both | 0 | 0 |
| **6049** | 1450 | 182 | 1 | 22368.387 | Coleraine Fm | 430 | | Coleraine | | | 90.825 | 95.022 | 30 | 0 | subsurface | 2 | 11 |
| **13347** | 3504 | 452 | 1 | 12938.724 | Tuscaloosa Fm | 2157 | | Tuscaloosa | | | 95.022 | 95.55 | 5 | 0 | surface | 0 | 0 |
| **13414** | 3534 | 454 | 1 | 10349.959 | Gordo Fm | 768 | | Gordo | Tuscaloosa | | 95.022 | 95.55 | 18 | 0 | both | 0 | 0 |
| **10702** | 2814 | 274 | 1 | 20481.164 | Kayak Shale of Endicott Gp | 3203 | | Kayak Shale | Endicott | | 345.5 | 357.05 | 300 | 0 | surface | 0 | 0 |
| **3934** | 778 | 101 | 1 | 18752.73 | Boice Shale | 4050 | | Boice Shale | | | 354.275 | 358.9 | 68 | 0 | subsurface | 0 | 0 |
| **4413** | 843 | 108 | 1 | 24896.395 | Boice Shale | 4050 | | Boice Shale | | | 357.05 | 358.9 | 3 | 0 | subsurface | 0 | 0 |
| **9101** | 2305 | 333 | 1 | 18458.613 | Gordon Shale | 4095 | | Gordon Shale | | | 506.6 | 509.35 | 91 | 42 | surface | 0 | 0 |
| **14024** | 3718 | 467 | 1 | 17942.957 | Carrara Fm | 324 | | Carrara | | | 509.35 | 515.726 | 450 | 0 | surface | 32 | 118 |
| **7599** | 2006 | 297 | 1 | 4416.283 | Harpers Fm | 840 | | Harpers | Chilhowee | | 517.255 | 525.039 | 900 | 370 | | 0 | 0 |
| **41907** | 11631 | 512 | 1 | 35648.656 | Roberts Draw | 73262 | | Roberts Draw | | | 1916.6675 | 1937.5 | 300 | 100 | | 0 | 0 |

<table>
<tr><th>lith</th><th>environ</th><th>econ</th><th>measure</th></tr>
<tr><td>ferruginous gravel siliciclastic sedimentary  ~ 0.1429|quartz sand siliciclastic sedimentary  ~ 0.7143|glauconitic sand siliciclastic sedimentary  ~ 0.1429</td><td>lacustrine indet. lacustrine non-marine |outwash plain glacial non-marine |estuary/bay fluvial marine</td><td>sand and gravel construction material</td><td></td></tr>
<tr><td>ferruginous quartz gravel siliciclastic sedimentary  ~ 0.1429|quartz sand siliciclastic sedimentary  ~ 0.7143|glauconitic sand siliciclastic sedimentary  ~ 0.1429</td><td>lacustrine indet. lacustrine non-marine |outwash plain glacial non-marine |estuary/bay fluvial marine</td><td>sand and gravel construction material</td><td></td></tr>
<tr><td>ferruginous quartz gravel siliciclastic sedimentary  ~ 0.1429|quartz sand siliciclastic sedimentary  ~ 0.7143|glauconitic sand siliciclastic sedimentary  ~ 0.1429</td><td>outwash plain glacial non-marine |estuary/bay fluvial marine |lacustrine indet. lacustrine non-marine</td><td>sand and gravel construction material</td><td></td></tr>
<tr><td>ferruginous quartz gravel siliciclastic sedimentary  ~ 0.1429|quartz sand siliciclastic sedimentary  ~ 0.7143|glauconitic sand siliciclastic sedimentary  ~ 0.1429</td><td>estuary/bay fluvial marine |lacustrine indet. lacustrine non-marine |outwash plain glacial non-marine</td><td>sand and gravel construction material</td><td></td></tr>
<tr><td>gravel siliciclastic sedimentary  ~ 0.4545|quartz sand siliciclastic sedimentary  ~ 0.4545|ferruginous chert chemical sedimentary  ~ 0.0909</td><td>fluvial indet. fluvial non-marine</td><td>sand and gravel construction material</td><td></td></tr>
<tr><td>ferruginous glauconitic limestone carbonate sedimentary  ~ 0.0909| mudstone siliciclastic sedimentary  ~ 0.4545|diatomaceous siltstone siliciclastic sedimentary  ~ 0.4545</td><td>deep subtidal shelf carbonate marine |shallow subtidal carbonate marine |coastal indet. siliciclastic marine</td><td></td><td></td></tr>
<tr><td>ferruginous glauconitic limestone carbonate sedimentary  ~ 0.0909| mudstone siliciclastic sedimentary  ~ 0.4545|diatomaceous siltstone siliciclastic sedimentary  ~ 0.4545</td><td>deep subtidal shelf carbonate marine |shallow subtidal carbonate marine |coastal indet. siliciclastic marine</td><td></td><td></td></tr>
<tr><td>conglomerate siliciclastic sedimentary  ~ 0.8333|ferruginous sandstone siliciclastic sedimentary  ~ 0.1667</td><td>marine  marine |non-marine  non-marine</td><td></td><td></td></tr>
<tr><td>siliciclastic siliciclastic sedimentary  ~ 0.7143| clay siliciclastic sedimentary  ~ 0.1429|ferruginous sand siliciclastic sedimentary  ~ 0.1429</td><td>non-marine  non-marine |fluvial indet. fluvial non-marine</td><td></td><td></td></tr>
<tr><td>clay siliciclastic sedimentary  ~ 0.1429|ferruginous sand siliciclastic sedimentary  ~ 0.1429|coarse fine siliciclastic siliciclastic sedimentary  ~ 0.7143</td><td>deltaic indet. siliciclastic marine |fluvial indet. fluvial non-marine |floodplain fluvial non-marine</td><td></td><td></td></tr>
<tr><td>ferruginous quartzite metasedimentary metamorphic  ~ 0.4545|black shale siliciclastic sedimentary  ~ 0.4545|ferruginous grainstone carbonate sedimentary  ~ 0.0909</td><td>marine  marine</td><td></td><td></td></tr>
<tr><td>greenish gray dolomitic shale siliciclastic sedimentary  ~ 0.7143| dolomite carbonate sedimentary  ~ 0.1429|red ferruginous shale siliciclastic sedimentary  ~ 0.1429</td><td>inferred marine  marine</td><td></td><td></td></tr>
<tr><td>greenish gray shale siliciclastic sedimentary  ~ 0.5556| dolomite carbonate sedimentary  ~ 0.1111|ferruginous oolitic grainstone carbonate sedimentary  ~ 0.1111|dolomitic silty shale siliciclastic sedimentary  ~ 0.1111|red sha</td><td>inferred marine  marine</td><td></td><td></td></tr>
<tr><td>green dark brown shale siliciclastic sedimentary  ~ 0.7143|ferruginous sandstone siliciclastic sedimentary  ~ 0.1429|glauconitic sandy greenish gray shale siliciclastic sedimentary  ~ 0.1429</td><td>inferred marine  marine</td><td></td><td></td></tr>
<tr><td>shale siliciclastic sedimentary  ~ 0.125| limestone carbonate sedimentary  ~ 0.625|glauconitic sandy shale siliciclastic sedimentary  ~ 0.125|ferruginous dolomite carbonate sedimentary  ~ 0.125</td><td>marine  marine</td><td></td><td></td></tr>
<tr><td>ferruginous sandstone siliciclastic sedimentary  ~ 0.3333| phyllite metasedimentary metamorphic  ~ 0.3333| arkose siliciclastic sedimentary  ~ 0.3333</td><td>inferred marine  marine</td><td></td><td></td></tr>
<tr><td>dolomite carbonate sedimentary  ~ 0.2778|sandy pelite siliciclastic sedimentary  ~ 0.2778| phyllite metasedimentary metamorphic  ~ 0.0556|ferruginous chert chemical sedimentary  ~ 0.2778| basalt volcanic igneous  ~ 0.055</td><td>inferred marine  marine</td><td></td><td></td></tr>
</table>

| notes | color | text_color | t_int_id | t_int_name | t_int_age | t_prop | units_above | b_int_id | b_int_name | b_int_age | b_prop | units_below | strat_name_long | refs | clat | clng | t_plat | t_plng | b_plat | b_plng |
|---|---|---|---|---|---|---|---|---|---|---|---|---|---|---|---|---|---|---|---|---|
| environ from EPA/USGS | #FCF768 | #000000 | 4 | Pleistocene | 0.0117 | 1 | 0 | 4 | Pleistocene | 2.58 | 0.77499 | 0 | Cape May Formation | 1 | 39.887 | -74.084 | 39.889 | -74.079 | 39.964 | -73.809 |
| environ from EPA/USGS | #FCF768 | #000000 | 4 | Pleistocene | 0.0117 | 0.83333 | 0 | 4 | Pleistocene | 2.58 | 0.66665 | 0 | Cape May Formation | 1 | 40.419 | -74.217 | 40.477 | -74.01 | 40.532 | -73.808 |
| environ from EPA/USGS | #FCF768 | #000000 | 4 | Pleistocene | 0.0117 | 0.83333 | 0 | 4 | Pleistocene | 2.58 | 0.66665 | 0 | Cape May Formation | 1 | 40.082 | -74.095 | 40.139 | -73.889 | 40.194 | -73.688 |
| environ from EPA/USGS | #FCF768 | #000000 | 4 | Pleistocene | 0.0117 | 0.83333 | 0 | 4 | Pleistocene | 2.58 | 0.66665 | 0 | Cape May Formation | 1 | 39.212 | -74.733 | 39.271 | -74.531 | 39.327 | -74.334 |
| environ- USGS and Stanford etal 2002 | #F58D2C | #000000 | 11 | Piacenzian | 2.58 | 1 | 0 | 11 | Piacenzian | 3.6 | 0.25 | 0 | Beacon Hill Gravel | 1 | 40.419 | -74.217 | 40.747 | -72.996 | 40.839 | -72.635 |
| environ- Miller & Aalto 2008 | #999999 | #000000 | 11 | Piacenzian | 2.58 | 0.1667 | 6694 | 12 | Zanclean | 5.333 | 0 | 0 | Pullen Formation | 1 | 40.557 | -124.349 | 41.329 | -122.812 | 41.753 | -121.94 |
| environ- Miller & Aalto 2008 | #999999 | #000000 | 11 | Piacenzian | 2.58 | 0.1667 | 6705 | 14 | Messinian | 7.246 | 0.74997 | 0 | Pullen Formation | 1 | 40.446 | -124.093 | 41.218 | -122.553 | 41.748 | -121.458 |
| environ- Bolin 1956 and Sloan 1964 | #FCF768 | #000000 | 39 | Turonian | 89.8 | 0.75 | 0 | 40 | Cenomanian | 100.5 | 0.83 | 0 | Coleraine Formation | 1 | 47.883 | -91.716 | 51.82 | -45.696 | 52.241 | -45.742 |
| environ- Marcher & Stearns 1962 | #A57B54 | #000000 | 40 | Cenomanian | 93.9 | 0.83 | 0 | 40 | Cenomanian | 100.5 | 0.75 | 0 | Tuscaloosa Formation | 1 | 36.03 | -86.753 | 39.867 | -44.419 | 39.882 | -44.258 |
| environ- Cahoon 1972 and Mancini et al 1987 | #A57B54 | #000000 | 40 | Cenomanian | 93.9 | 0.83 | 0 | 40 | Cenomanian | 100.5 | 0.75 | 0 | Gordo Formation | 1 | 34.963 | -87.379 | 38.927 | -45.346 | 38.938 | -45.18 |
| | #999999 | #000000 | 144 | Osagean | 343.5 | 0.75 | 10701 | 145 | Kinderhookian | 358.9 | 0.25 | 10703 | Kayak Shale | 1 | 67.048 | -150.095 | 9.358 | -24.946 | 7.219 | -29.847 |
| | #999999 | #000000 | 145 | Kinderhookian | 351.5 | 0.625 | 3933 | 145 | Kinderhookian | 358.9 | 0 | 0 | Boice Shale | 1 | 39.446 | -95.773 | -26.409 | -56.791 | -27.315 | -57.217 |
| | #999999 | #000000 | 145 | Kinderhookian | 351.5 | 0.25 | 0 | 145 | Kinderhookian | 358.9 | 0 | 0 | Boice Shale | 1 | 37.766 | -96.751 | -27.081 | -59.24 | -27.447 | -59.29 |
| | #999999 | #000000 | 128 | Middle Cambrian | 501 | 0.44 | 9100 | 128 | Middle Cambrian | 511 | 0.165 | 9102 | Gordon Shale | 1 | 48.181 | -112.762 | 34.401 | -121.851 | 33.954 | -124.662 |
| | #4D52E7 | #E0E0E0 | 128 | Middle Cambrian | 501 | 0.165 | 14022 | 130 | Early Cambrian | 538.8 | 0.83 | 14025 | Carrara Formation | 1 | 36.42 | -115.434 | 30.815 | -138.296 | 26.293 | -144.692 |
| S.S. Desc: https://pubs.usgs.gov/bul/2123/report.pdf<br>-lower: meta-arkose<br>-upper: ferruginous, magnetite rich s.s. | #5E3006 | #000000 | 130 | Early Cambrian | 511 | 0.775 | 7598 | 130 | Early Cambrian | 538.8 | 0.495 | 7600 | Harpers Formation | 1 | 39.696 | -77.995 | 1.424 | -126.867 | -14.696 | -138.832 |
| | #4D52E7 | #E0E0E0 | 270 | Orosirian | 1800 | 0.53333 | 41908 | 270 | Orosirian | 2050 | 0.45 | 41905 | Roberts Draw Formation | 1 | 43.949 | -104.206 | 0 | 0 | 0 | 0 |
